# Mapping the neutralizing specificity of human anti-HIV serum by deep mutational scanning

**DOI:** 10.1101/2023.03.23.533993

**Authors:** Caelan E. Radford, Philipp Schommers, Lutz Gieselmann, Katharine H. D. Crawford, Bernadeta Dadonaite, Timothy C. Yu, Adam S. Dingens, Julie Overbaugh, Florian Klein, Jesse D. Bloom

## Abstract

Understanding the specificities of human serum antibodies that broadly neutralize HIV can inform prevention and treatment strategies. Here we describe a deep mutational scanning system that can measure the effects of combinations of mutations to HIV envelope (Env) on neutralization by antibodies and polyclonal serum. We first show that this system can accurately map how all functionally tolerated mutations to Env affect neutralization by monoclonal antibodies. We then comprehensively map Env mutations that affect neutralization by a set of human polyclonal sera known to target the CD4-binding site that neutralize diverse strains of HIV. The neutralizing activities of these sera target different epitopes, with most sera having specificities reminiscent of individual characterized monoclonal antibodies, but one sera targeting two epitopes within the CD4 binding site. Mapping the specificity of the neutralizing activity in polyclonal human serum will aid in assessing anti-HIV immune responses to inform prevention strategies.

## Introduction

Efforts to create a HIV vaccine have been stymied in part by rapid and continuing diversification of the virus’s envelope (Env) protein^1, 2^. However, some individuals with HIV do naturally develop polyclonal serum antibody responses to Env that broadly neutralize many viral strains^3–5^. Much progress has been made characterizing individual broadly neutralizing antibodies. However, individual antibodies do not always recapitulate the neutralizing activity of the serum of the individuals from whom they were isolated^6–9^.

Mapping the specificity of polyclonal neutralizing serum antibodies is more difficult than characterizing individual monoclonal antibodies. One important advance has been the development of electron microscopy-based polyclonal epitope mapping (emPEM) methods to visualize how multiple different serum antibody Fabs bind to Env^10–12^. However, this approach characterizes binding rather than neutralizing specificity, and one major finding from emPEM is that many serum antibodies bind non-neutralizing epitopes^10–13^. Fingerprinting approaches can define neutralizing epitopes, but does not provide mutation-level specificity and requires making measurements for large virus panels^14, 15^. Deep mutational scanning can map Env mutations that escape antibody neutralization^13, 16–19^. However, existing HIV deep mutational scanning work has used approaches that are only able to look at effects of individual mutations, which is a limitation when trying to map polyclonal serum antibodies that may target multiple epitopes^13^.

Precisely mapping neutralizing specificities and escape mutations is especially challenging for antibodies that target the CD4-binding site. Such antibodies recognize conserved Env residues while typically avoiding steric clashes with glycans rather than depending on them for neutralization, unlike antibodies targeting other epitopes such as the V1/V2 loops or V3 loop^3, 4^. As a result, CD4-binding site targeting antibodies can have near pan-HIV neutralization breadth and high potency despite sequence and glycan heterogeneity across strains of HIV^3–5^, and are therefore promising candidates for treatment and prophylaxis strategies^5, 20, 21^. But the higher conservation of their epitopes can also make it more difficult to map escape mutations for such antibodies^17^.

Here we use an improved deep mutational scanning system to measure how mutations affect neutralization of Env by human anti-HIV sera that target the CD4-binding site. This new system can measure the effects of combinations of mutations, enabling quantitative deconvolution of how mutations mediate escape at distinct antibody epitopes^22^. We find that several sera have neutralizing activities that resemble monoclonal antibodies, but one sera has neutralizing activity targeting two distinct epitopes. These maps shed light on the specificity of human serum that can broadly neutralize many HIV strains. In addition, the method we employ could be used in the future to evaluate and compare the neutralizing specificities of anti-HIV sera elicited by different vaccine regimens.

## Results

### Single-round replicative lentivirus deep mutational scanning platform for HIV Env

We recently described a deep mutational scanning platform based on a single-round replicative lentivirus that does not encode any viral genes except for the viral entry protein^23^, which in our current study is Env. This platform enables creation of large libraries of single-round replicative lentiviruses with a genotype-phenotype link between barcodes in the lentivirus genomes and the entry proteins on the surfaces of virions (Figure 1A,B). Key aspects of this platform include encoding viral entry protein mutants in lentivirus genomes with random nucleotide barcodes and using a lentivirus genome with a full 3’ LTR that can be reactivated after infection^1–3^ (Figure 1A). Creation of the mutant libraries involves a two-step process of first integrating lentivirus genomes into cells at just one copy per cell, and then generating mutant virus libraries with a genotype-phenotype link (Figure 1B). PacBio sequencing is used to link Env mutants with their nucleotide barcodes, and later experiments use short read Illumina sequencing of the barcodes to measure mutant frequencies.

**Figure 1:**
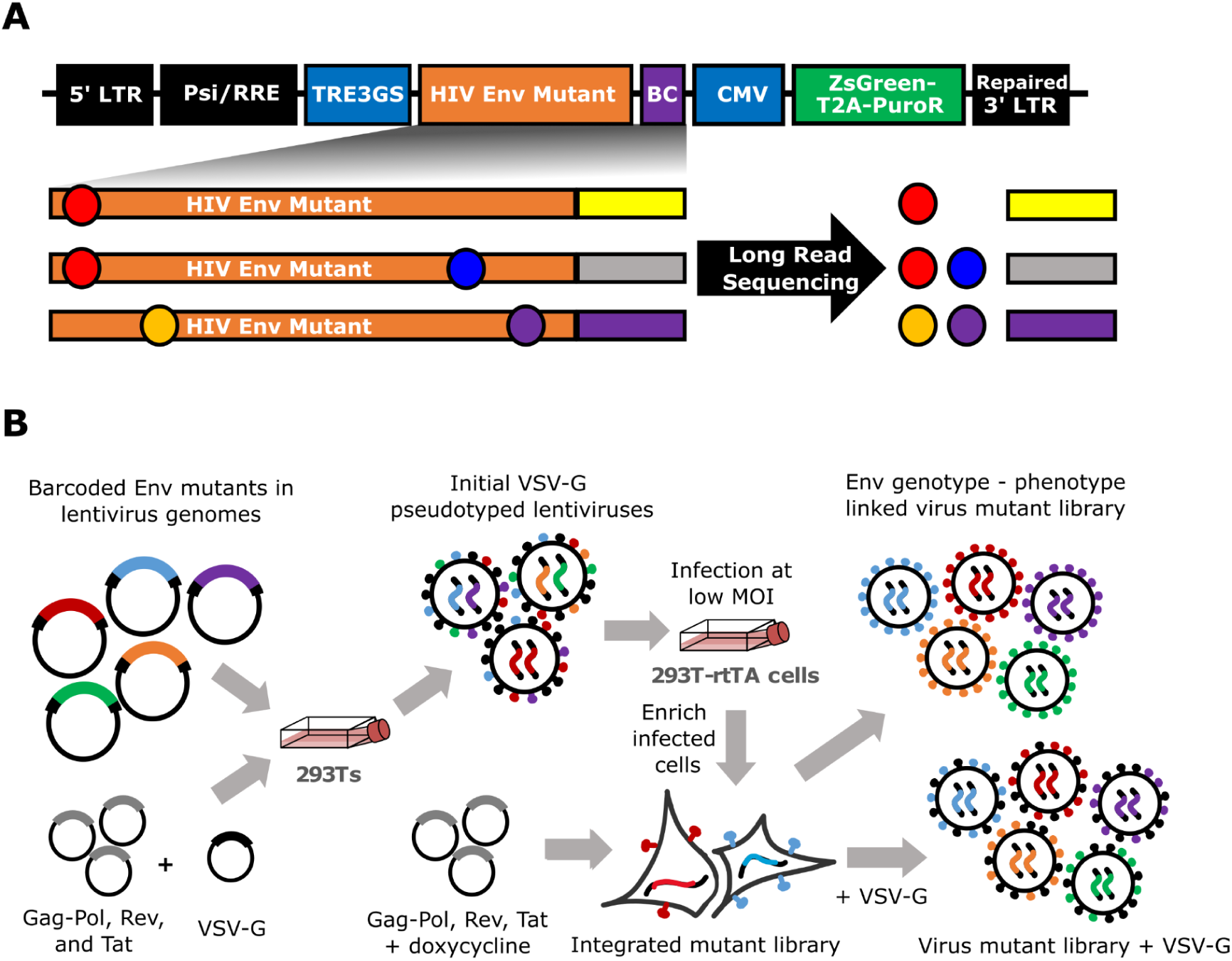
Lentivirus platform for deep mutational scanning. (A) The lentivirus genome used for deep mutational scanning. The genome contains the full 5’ and 3’ LTR sequences, including the U3 sequence usually deleted in the 3’ LTR. Env is under control of an inducible TRE3G promoter and followed by a 16N random nucleotide barcode. A CMV promoter drives ZsGreen and puromycin resistance (PuR) expression. (B) Approach for generating genotype-phenotype linked variant libraries. Lentivirus genomes carrying barcoded Env mutants are transfected into 293T cells alongside plasmids expressing the lentiviral proteins necessary for creating single-cycle infectious virions and VSV-G. The resulting VSV-G pseudotyped viruses are used to infect 293T-rtTA cells at a low multiplicity of infection, such that most infected cells receive just one viral genome. Infected cells are enriched via puromycin selection, and genotype-phenotype linked Env-expressing virus variant libraries are generated by inducing Env expression with doxycycline and transfecting plasmids encoding the lentivirus genes. The virus variant libraries are also generated separately with VSV-G, and these VSV-G pseudotyped viruses can infect cells regardless of whether or not they have a functional Env and so can be used to readout the library composition.

### Env mutant library design and generation

Our libraries used Env from the transmitted/founder virus BF520.W14M.C2^26, 27^ (referred to hereafter as BF520). We chose this Env for several reasons. First, since it is from a transmitted/founder virus, it is particularly relevant for antibody neutralization studies^6, 27, 28^. Second, the BF520 Env yielded high titers in our lentiviral deep mutational scanning platform (Supplemental Figure 1). Third, our lab has previously performed deep mutational scanning of the BF520 Env using full-length replicative HIV libraries in a prior system that did not enable analysis of multiply mutated Env mutants^16, 29^, thereby providing comparator data to benchmark the current study.

A goal of our experiments is to map escape from polyclonal serum antibodies as well as monoclonal antibodies. Since polyclonal serum is composed of multiple antibodies that can target different epitopes^30–32^, mapping escape from serum requires libraries that contain Envs with mutations in multiple epitopes^22^. However, about half of all possible amino-acid mutations to proteins are highly deleterious,^29, 33–35^ so a library of multiply mutated Envs with random mutations would contain a high fraction of non-functional proteins. Therefore, we designed the libraries to exclude most highly deleterious mutations. We also mutagenized only the Env ectodomain (and not the signal peptide, transmembrane domain, and cytoplasmic tail), since neutralizing antibodies always bind the ectodomain.

To design libraries containing mostly functional Env mutants, we drew on two sources of information. The first source of information was prior deep mutational scanning data for BF520 Env generated using full-length replicative HIV virions in a system that could only measure the average effect of mutations across different genetic backgrounds^16, 29^. We used data from this prior deep mutational scanning to identify well-tolerated mutations (Figure 2A, left panel). The second source of information was an alignment of group M HIV-1 sequences^36^, which we used to identify any mutations relative to BF520 present more than once in natural sequences (Figure 2A, middle panel). Our library design included the 7110 amino-acid mutations in the BF520 ectodomain that were either tolerated in the prior deep mutational scanning or present multiple times in the natural sequence alignment (Figure 2A, right panel). We created these Env libraries using a previously described PCR mutagenesis approach modified to target these specific mutations^23, 37^.

**Figure 2:**
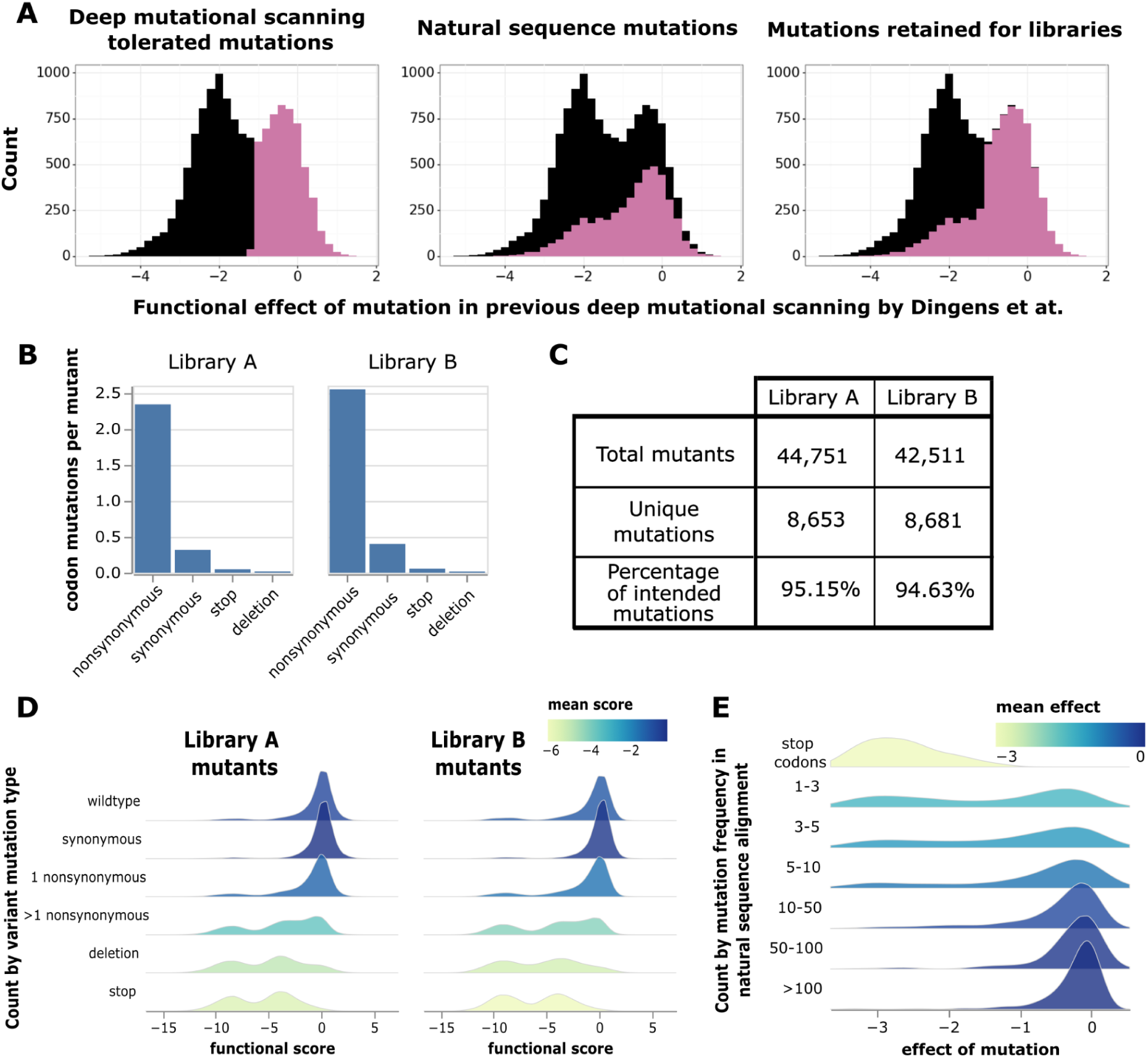
Mutant library design and functional effects of mutations. (A) Choice of targeted mutations based on measured effects in prior deep mutational scanning^29^ and occurrences in natural HIV sequences. The distributions of previously measured mutation effects are shown for all mutations to BF520 (black) with highlighting of subsets of mutations (purple). From left to right, highlighted are mutations well tolerated in the prior deep mutational scanning^29^, mutations found multiple times in natural sequences, and the union of these two sets. Mutations in the union of the two sets were designed into our libraries. (B) Average codon mutations per Env mutant, separated by type of codon mutation. (C) Total number of barcoded Env mutants in each library, along with the numbers of unique mutations and percentage of the intended mutations present across these mutants. (D) Distributions of functional scores measured in our deep mutational scanning across Env mutants, separated by the types of codon mutations found in the mutants. Negative functional scores indicate impaired Env-mediated infection relative to unmutated BF520 Env. Histograms are colored by mean functional score. (E) Distributions of mutation effects versus how often that mutation is found in natural sequences. The distribution of stop codon effects is also shown.

We generated two independent Env mutant libraries to perform the deep mutational scanning in biological duplicate. PacBio sequencing showed that each library had ∼2.5 nonsynonymous mutations per Env mutant, which are linked via the barcode and so can be evaluated in combination (Figure 1A). There was a low frequency of synonymous mutations, stop codons, and in-frame deletions (Figure 2B). Overall, ∼84% of the mutations were among the 7,110 mutations targeted in our library design. Each library contained ∼40,000 barcoded mutants, and together the two libraries sampled ∼97% of the targeted mutations (Figure 2C).

### Effects of mutations on Env-mediated viral entry

To measure how mutations affected the ability of Env to mediate viral infection in cell culture, we generated libraries pseudotyped with just the Env mutants or also pseudotyped with VSV-G (Figure 1B). We then used these libraries to infect TZM-bl cells^38–40^, which express Env’s primary receptor (CD4) and co-receptors (CCR5 and CXCR4). We sequenced the barcodes of virions that infected cells in each condition: all virions are expected to infect cells when VSV-G is present, but only virions with functional Envs will infect cells in the absence of VSV-G. Each barcoded Env variant was assigned a functional score calculated as the log of the ratio of the frequency of that variant (relative to unmutated BF520 Env) in the Env versus VSV-G mediated infections. Negative functional scores indicate an Env mutant is worse at infecting cells than unmutated BF520 Env, while positive functional scores indicate it is better at infecting cells.

As expected, Env mutants with only synonymous mutations have “wildtype-like” functional scores of near zero, while mutants with stop codons generally have highly negative functional scores (Figure 2D). Most mutants in the libraries with only one nonsynonymous mutation have functional scores close to zero, suggesting our library design largely incorporated functionally tolerated mutations as intended. Env mutants with multiple nonsynonymous mutations more often have substantially negative functional scores, as expected from the accumulation of multiple sometimes deleterious mutations (Figure 2D).

To estimate the effect of each individual mutation on viral entry, we fit global epistasis models to the functional scores^41, 42^. The effects of mutations on Env-mediated viral entry are shown at https://dms-vep.github.io/HIV_Envelope_BF520_DMS_CD4bs_sera/muteffects_observed_heatmap.html in an interactive heatmap. Mutations found more often among natural sequences tend to have more favorable effects in our experiments than mutations only found rarely among natural sequences (Figure 2E), suggesting mutations that are favorable for viral entry in our experiments are generally also favorable during natural HIV evolution.

### Accurate mapping of effects of Env mutations on antibody neutralization

We next used the deep mutational scanning platform to map how Env mutations affect antibody neutralization. As a first proof-of-principle, we mapped mutational escape from neutralization by the well-characterized broadly neutralizing antibody PGT151^43^. To directly quantify how mutations affected neutralization, we included a non-neutralized “standard” virus pseudotyped with VSV-G to enable conversion of relative sequencing counts to absolute neutralization measurements^23^. To estimate the effects of individual mutations from our library measurements (which include both singly and multiply mutated Envs), we fit a biophysical model where antibody neutralization at each epitope has a Hill-curve dependence on antibody concentration and mutations within a given epitope have additive effects on antibody affinity^22^. This model, which is implemented in the *polyclonal* software (https://jbloomlab.github.io/polyclonal/), utilizes information from both singly and multiply mutated Env variants under realistic assumptions about how mutations combine to escape antibody binding.

Our mapping showed that PGT151 is escaped by mutations in the fusion peptide or affecting N-linked glycans recognized by PGT151 (Figure 3A,B and interactive escape maps linked in figure legend). In particular, PGT151 is strongly escaped by any mutations knocking out the N611 glycan, specific mutations at the N637 glycan, mutations to positively charged residues at sites around the N637 glycan, mutations at sites 647 and 648, and mutations at sites 512 and 514 in the fusion peptide (Figure 3A,B). We also mapped lower magnitude escape at sites 537-543. All these mutations are in or near the binding footprint of PGT151^44^ (Figure 3C).

**Figure 3:**
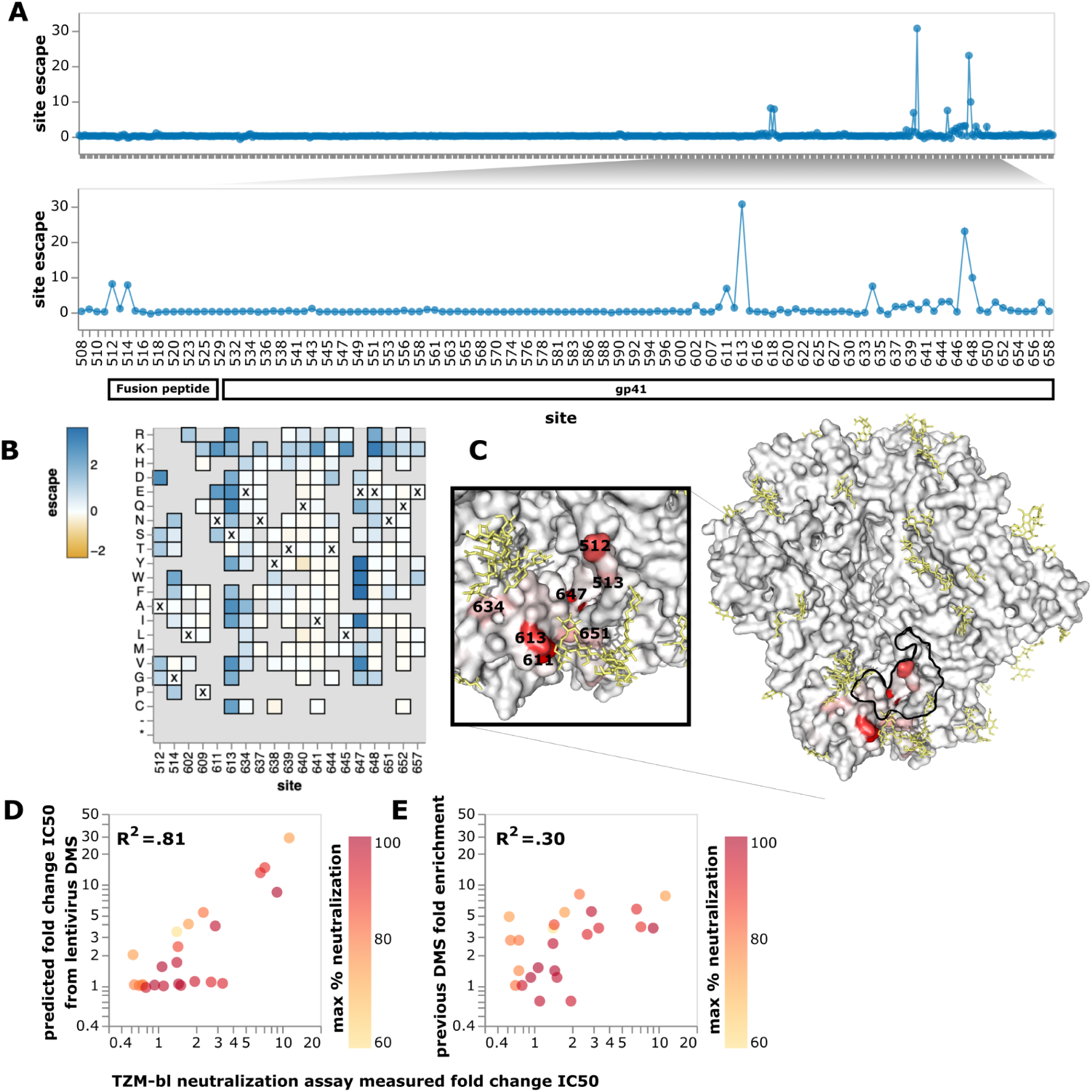
Neutralization escape map for antibody PGT151. (A) The top panel shows PGT151 escape across all sites in the BF520 Env ectodomain, and the bottom panel zooms into key sites. The y-axis shows escape summed across all mutations at each site. (B) Heatmap of effects of individual mutations at key sites of escape. Residues marked with X are wildtype residues in BF520. Residues grayed out are not present in the variant libraries, typically because they are deleterious for Env function. See https://dms-vep.github.io/HIV_Envelope_BF520_DMS_CD4bs_sera/PGT151_escape_plot.html for an interactive version of the site and mutation escape plots. (C) Site escape mapped onto a structure of PGT151-bound Env, with red indicating sites where mutations cause escape. Residues within 4 angstroms of antibody PGT151 in the structure are outlined in black. Glycans are colored yellow. This visualization was generated using the structure of BG505 ΔCT N332T (PDB 6MAR, antibody PGT151 removed). (D) Correlation of predicted fold-change in IC50 from the current deep mutational scanning versus fold change in IC50 as previously measured in TZM-bl neutralization assays^16^. Mutations are colored by the max neutralization plateau observed for that variant in TZM-bl neutralization assays using PGT151. Pearson correlation coefficients were calculated on the fold-changes on a linear scale. Note the fold change IC50 predicted by deep mutational scanning is calculated using the full biophysical model fit to the data (see Methods). (E) Correlation of fold enrichment in prior HIV Env deep mutational scanning^16^ versus fold-change in IC50 for the same set of single mutants.

The escape predicted by our deep mutational scanning was highly correlated with IC50s measured in previously performed TZM-bl neutralization assays^16^ (Figure 3D). The correlation with neutralization IC50s was substantially better for our current deep mutational scanning than an earlier approach that used libraries of HIV virions in a system where it was not possible to measure multiple mutations or absolute neutralization^16^ (Figure 3D,E).

### Broadly neutralizing anti-HIV sera

We assembled a set of sera from individuals with HIV to test if we could map neutralizing specificities in a polyclonal context^17^. We chose sera based on their ability to broadly neutralize a global HIV panel^45^ and potently neutralize BF520 pseudovirus. Based on these criteria, we chose four sera collected from individuals in Germany living with HIV (Supplemental Figure 2A,B). Based on the f61 neutralization fingerprinting panel^14^, these sera were predicted to be primarily VRC01-like, meaning they target the CD4-binding site (Supplemental Figure 2C). Note that all the sera in our study target the CD4-binding site because we chose broad sera that neutralized BF520 (which is relatively resistant to V3 antibodies^6^); neutralizing human anti-HIV sera can target other epitopes^14, 46, 47^. Importantly, we used purified IgGs from these sera for our experiments, since antiretroviral drugs present in the sera could interfere with our lentiviral-based assays.

### Neutralization escape maps of serum IDC561 and its constituent antibody 1-18 are similar

We first analyzed serum IDC561, which was collected from the same individual from whom the broadly neutralizing antibody 1-18 was isolated^17^. The antibody was isolated from B cells from the same blood draw date as the serum we used, suggesting antibody 1-18 is likely present in the serum. We mapped escape from neutralization by antibody 1-18 alongside the serum in order to compare the escape maps. It has been previously reported that 1-18 and purified IgGs from IDC561 display similar neutralization of a panel of viral strains and mutants, suggesting that neutralization by serum IDC561 is dominated by 1-18^17^. We therefore hypothesized that the neutralization escape map of serum IDC561 might resemble that of 1-18.

The maps for serum IDC561 and antibody 1-18 generally show neutralization escape at the same sites in Env, although the relative magnitude differs between the serum and antibody (Figure 4, and interactive escape maps linked in figure legend). In particular, both the serum and antibody are escaped by mutations around the V1/V2 loop, at β20/β21, and at the β23-V5-β24 structure (Figure 4A,B,C,D). Around the V1/V2 loop, the greatest escape from 1-18 is by mutations at site 198 in the middle of the N197 glycosylation motif (Figure 4A,B,D), and by mutations to sites 202, 203, and 206 (Figure 4A,B,D). IDC561 is also escaped by mutations at site 198, but mutations at sites 202 and 203 cause more escape for the serum than for 1-18, while there is less escape at site 206 for the serum than for 1-18 (Figure 4A,C,D). At β20/β21, mutations at sites 428-430 escape both 1-18 and IDC561, but the magnitude of this effect is lower for IDC561 than 1-18 (Figure 4A,B,C,D). At the β23-V5-β24 structure, mutations to sites 471, 474 and 476 escape 1-18, but only mutations at site 471 strongly escape IDC561 (Figure 4A,B,C,D).

**Figure 4:**
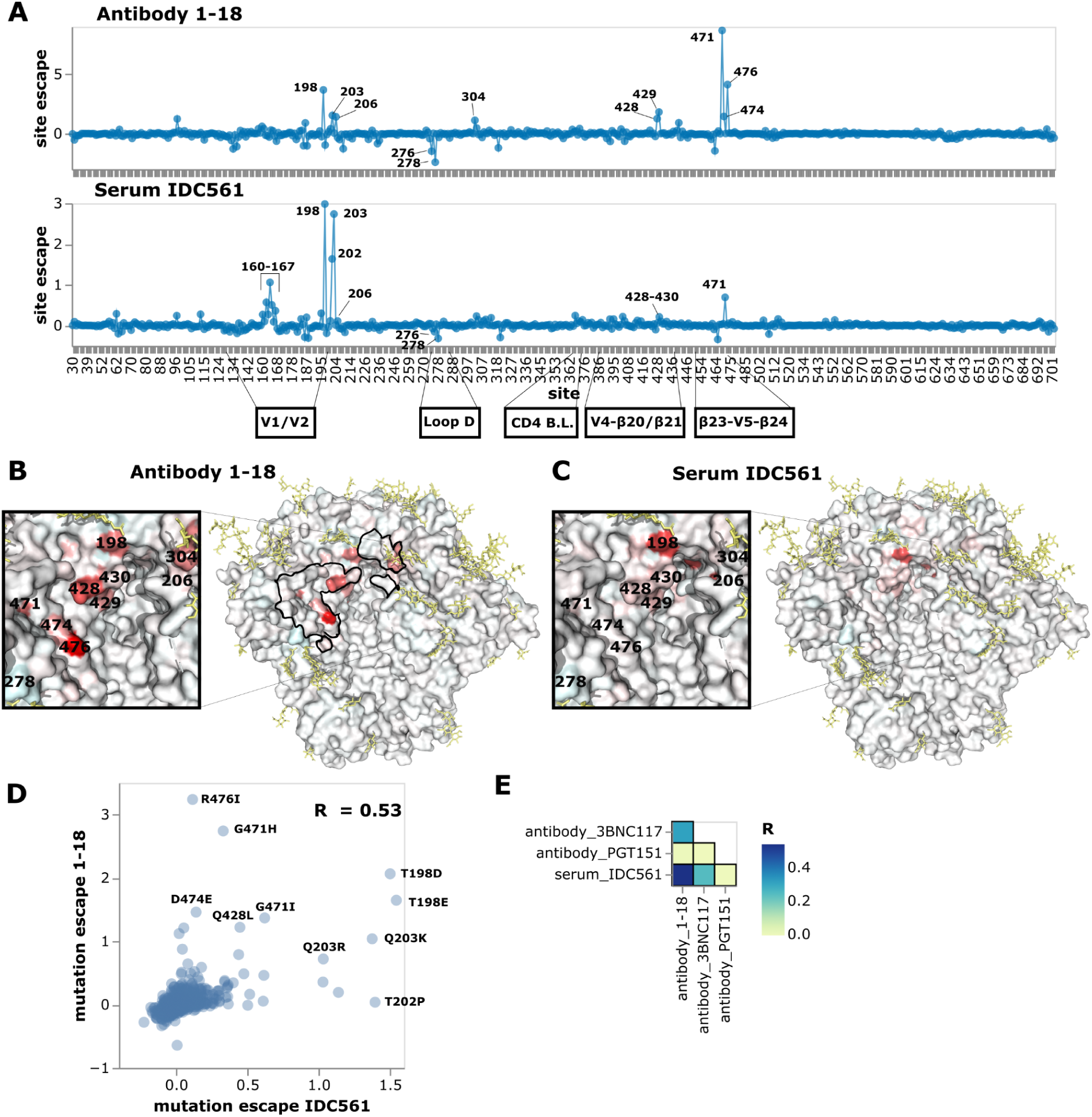
Neutralization escape map for antibody 1-18 and purified IgGs from IDC561. **(A)** Escape at all sites in the BF520 Env ectodomain from antibody 1-18 and serum IDC561. Positive values represent sites where mutations cause escape on average, while negative values represent sites where mutations enhance neutralization on average. See https://dms-vep.github.io/HIV_Envelope_BF520_DMS_CD4bs_sera/1-18_escape_plot.html and https://dms-vep.github.io/HIV_Envelope_BF520_DMS_CD4bs_sera/IDC561_escape_plot.html for interactive versions of the escape maps for 1-18 and IDC561, respectively. **(B)** Site escape from 1-18 mapped onto a structure of 1-18-bound Env. Residues within 4 angstroms of antibody 1-18 in the structure are outlined in black. This visualization was generated using the structure of BG505.SOSIP.664 (PDB 6UDJ, antibodies 10-1074 and 1-18 removed)^17^. (C) Site escape from IDC561 mapped onto the same structure. (D) Scatter plot of how mutations escape serum IDC561 versus antibody 1-18. (E) Correlations of how mutations escape IDC561 versus antibodies 1-18, 3BNC117, or PGT151.

The escape map for serum IDC561 was substantially more similar to that of antibody 1-18 than another CD4 binding site antibody, 3BNC117^48^, as well as the fusion peptide/gp120-gp41 interface-targeting antibody PGT151 (Figure 4E). This similarity suggests that antibody 1-18, which was isolated from the individual from which serum IDC561 was obtained, contributes substantially to overall neutralization by this serum as suggested by prior studies^17^. However, the fact that the serum IDC561 map does not entirely mirror that of 1-18 shows that other antibodies or members of the same clonal family also contribute to serum neutralization.

### Escape maps of other sera show diverse patterns of neutralization specificity

We next analyzed three more CD4-binding site targeting sera. The first of these sera, IDC513, was most escaped by mutations in loop D, similar to the well-characterized antibody 3BNC117 (Figure 5, and interactive escape maps linked in figure legend), although that antibody was not isolated from this individual. Both 3BNC117 and IDC513 are escaped by mutations in loop D, particularly at site 281 (Figure 5A-D). However, mutations at sites 276 and 278, which knockout the N276 glycan, enhance neutralization by both 3BNC117 and IDC513 (Figure 5A-D). These mutations also sensitize Env to neutralization by 1-18 and IDC561, but not to the same extent (Figure 4A). Mutations at sites 456, 459, and 471 in the β23-V5-β24 structure also escape both IDC513 and 3BNC117, and there is lower magnitude escape by mutations in and around the CD4 binding loop and other variable loops (Figure 5A-C). Overall the escape map for IDC513 correlates better with 3BNC117 than 1-18 (Figure 5D,E). Because 3BNC117 and serum IDC513 are from different individuals, neutralizing antibodies in serum IDC513 must have convergently evolved to target similar sites as antibody 3BNC117. Convergent evolution of broadly neutralizing HIV antibodies from the same heavy chain genes has been observed previously^48^, although we do not know the genes encoding the neutralizing antibodies in serum IDC513. Note that efforts to induce similar antibody specificities form the basis of some vaccine strategies^49, 50^.

**Figure 5:**
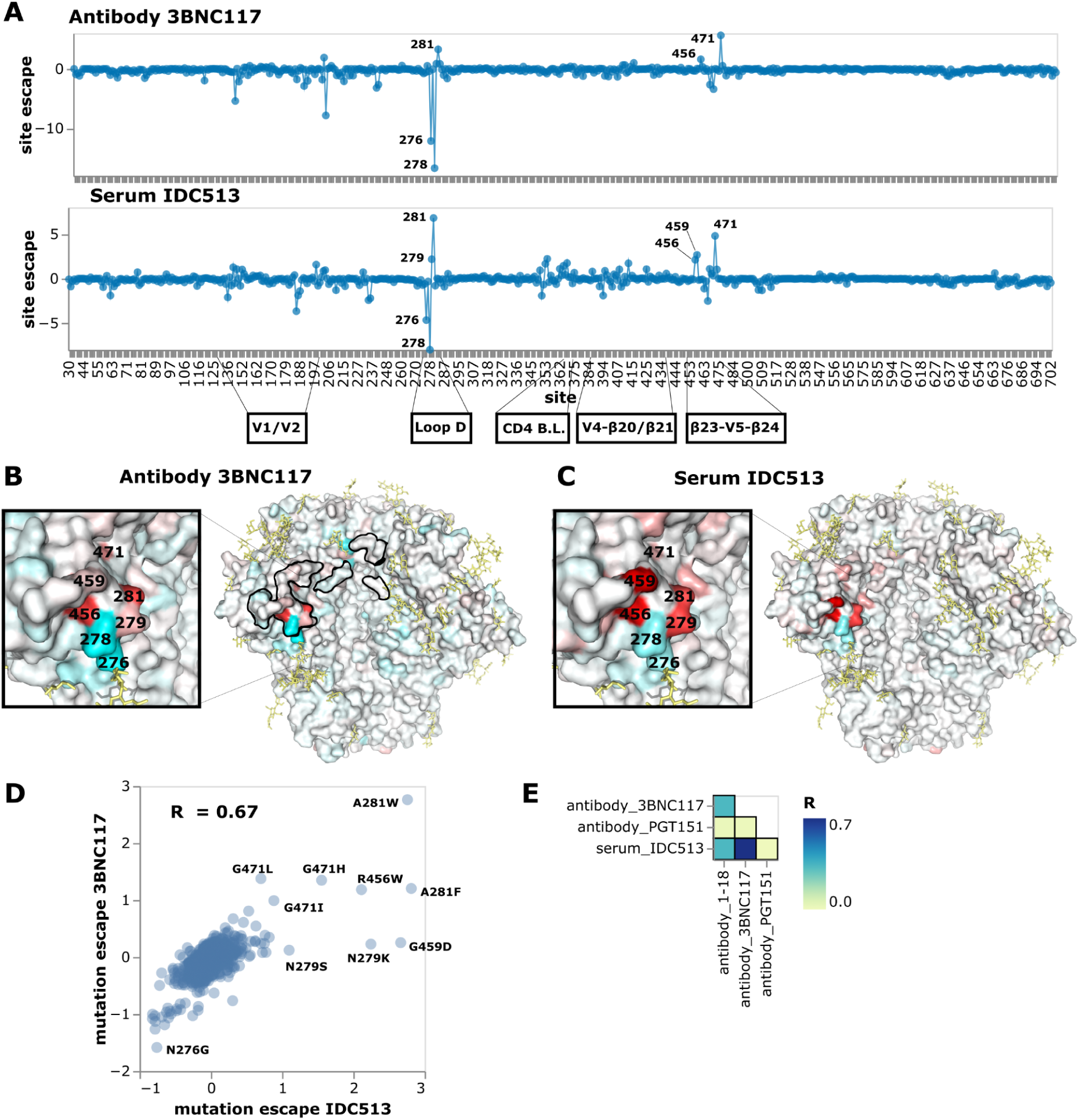
Neutralization escape maps for antibody 3BNC117 and purified IgGs from IDC513. **(A)** Escape at all sites in BF520 Env ectodomain from antibody 3BNC117 and serum IDC513. See https://dms-vep.github.io/HIV_Envelope_BF520_DMS_CD4bs_sera/3BNC117_escape_plot.html and https://dms-vep.github.io/HIV_Envelope_BF520_DMS_CD4bs_sera/IDC513_escape_plot.html for interactive versions of the escape maps for 3BNC117 and IDC513, respectively. **(B)** Site escape from 3BNC117 mapped onto a structure of 3BNC117-bound Env. Residues within 4 angstroms of antibody 3BNC117 in the structure are outlined in black. This visualization was generated using the structure of BG505.SOSIP.664 along with antibody 3BNC117 (PDB 5V8M)^55^. (C) Site escape from IDC513 mapped onto the same structure. (D) Scatter plot of how mutations escape serum IDC513 versus antibody 3BNC117. (E) Correlations of how mutations escape IDC513 versus antibodies 1-18, 3BNC117, or PGT151.

In contrast to IDC513 and IDC561, the escape map of IDF033 reveals a dependence on the N276 glycan for neutralization (Figure 6A,B, and interactive escape maps linked in the figure legend). Mutations at sites 276 and 278 that ablate the N276 glycan cause by far the greatest escape (Figure 6A,B). Other mutations in loop D, particularly at site 281, also more weakly escape from IDF033 (Figure 6A,B). At the β23-V5-β24 structure, mutations at sites 463 and 465 of the N463 glycosylation motif enhance neutralization by IDF033, but the mutation N463S causes escape by shifting the glycosylation motif to N461 (Figure 6A, Supplemental figure 3A). Other nearby sites also have mutation-specific effects (Supplemental figure 3A). For example at site S460, only some of the amino-acid changes cause escape (Supplemental figure 3A). Note that the neutralization fingerprinting panel (Supplementary Figure 2C) suggests serum IDF033 also has some V3-targeting activity, but this is not apparent in our escape maps probably because BF520 has a relatively high baseline resistance to V3 targeting antibodies^6^.

**Figure 6:**
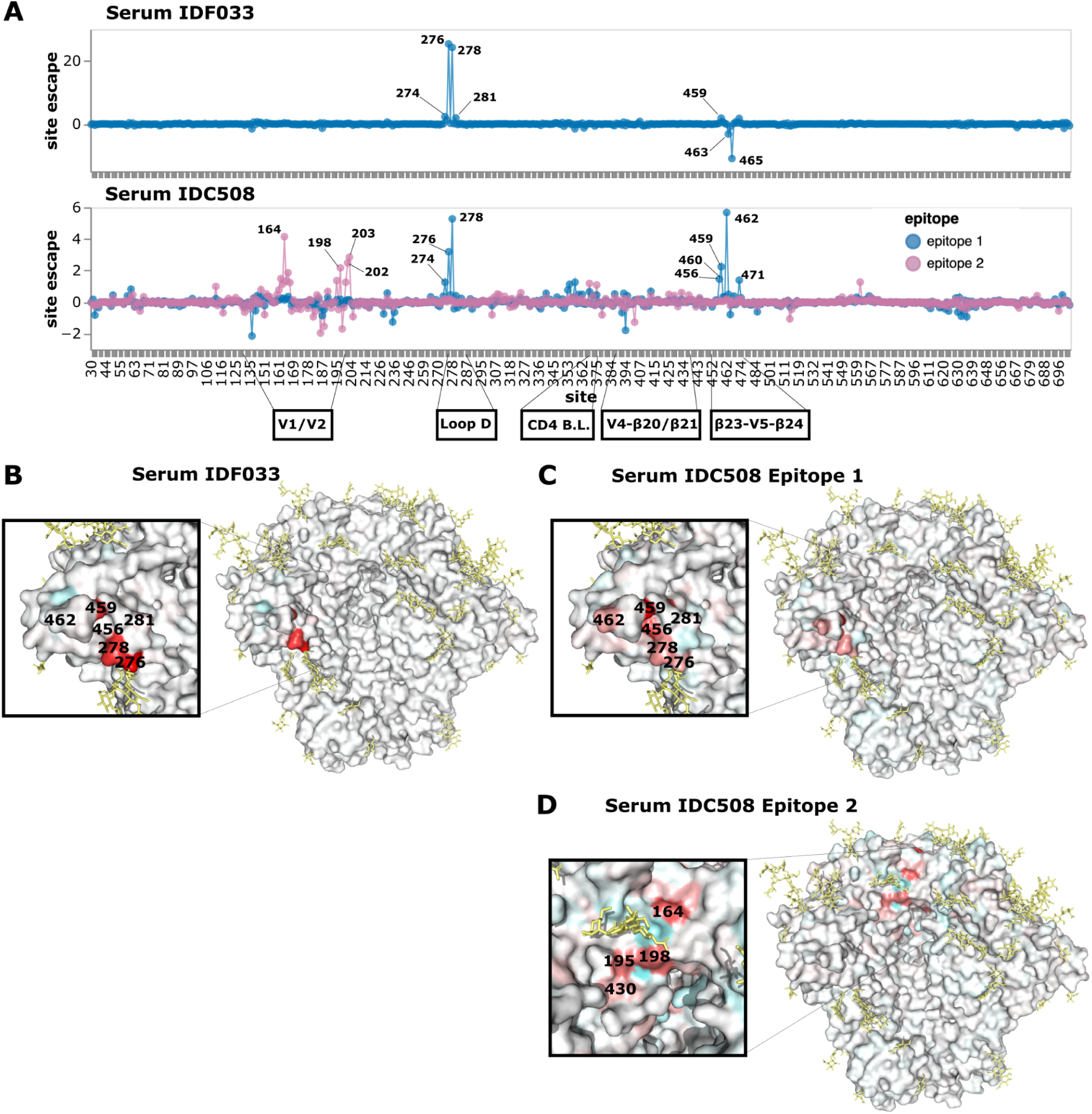
Neutralization escape maps for purified IgGs from IDF033 and IDC508. (A) Escape at all sites in BF520 Env ectodomain from serum IDF033 and serum IDC508. See https://dms-vep.github.io/HIV_Envelope_BF520_DMS_CD4bs_sera/IDF033_escape_plot.html and https://dms-vep.github.io/HIV_Envelope_BF520_DMS_CD4bs_sera/IDC508_escape_plot.html for interactive versions of the escape maps for IDF033 and IDC508, respectively. (B) Site escape from IDF033 mapped onto a structure of Env. This visualization was generated using the structure of BG505.SOSIP.664 (PDB 6UDJ, antibodies 10-1074 and 1-18 removed)^17^. (C) Site escape from the first IDC508 epitope mapped onto the same structure. (D) Site escape from the second IDC508 epitope mapped onto the same structure.

The escape map for the final serum, IDC508, revealed neutralization escape at two distinct antibody epitopes (Figure 6A,C,D, and interactive escape maps linked in figure legend). The existence of two epitopes was inferred by fitting the biophysical model^22^ to the deep mutational scanning measurements and finding that escape in multiply mutated variants was best explained by mutations affecting antibody binding at two distinct regions. Note that identification of two separate epitopes is crucially enabled by the ability of our deep mutational scanning system to quantify escape by Envs with multiple mutations^22^.

The first IDC508 epitope depends on the presence of the N276 glycan for neutralization and therefore is escaped by mutations at sites 276 and 278 as well as other mutations in loop D, similar to IDF033 (Figure 6A,C). Neutralization at this first epitope is also escaped by mutations at the β23-V5-β24 structure, also similar to IDF033 (Figure 6A,B,C, Supplemental Figure 3B). The second IDC508 epitope mapped mainly to sites around the V1/V2 loop (Figure 6A,D). Mutations at site 198 cause escape from neutralization at this second epitope, similar to 1-18 and IDC561 (Figure 4A,C,D, Figure 6A,D). Mutations at site 201, 202, and 203 and in the V2 loop at sites 160-167 also escape at the second epitope, again similar to IDC561 (Figure 4A, Figure 6A,D). Therefore, each of the two epitopes targeted by the neutralizing activity of IDC508 resembled the epitope targeted by another serum.

### Deep mutational scanning escape maps validate in neutralization assays

We validated the deep mutational scanning by performing pseudovirus neutralization assays on single amino-acid mutants of Env with a range of effects in the escape maps. The changes in pseudovirus neutralization assay IC80’s correlated well with the mutational effects predicted by the deep mutational scanning for all four sera (Figure 7A,B).

**Figure 7:**
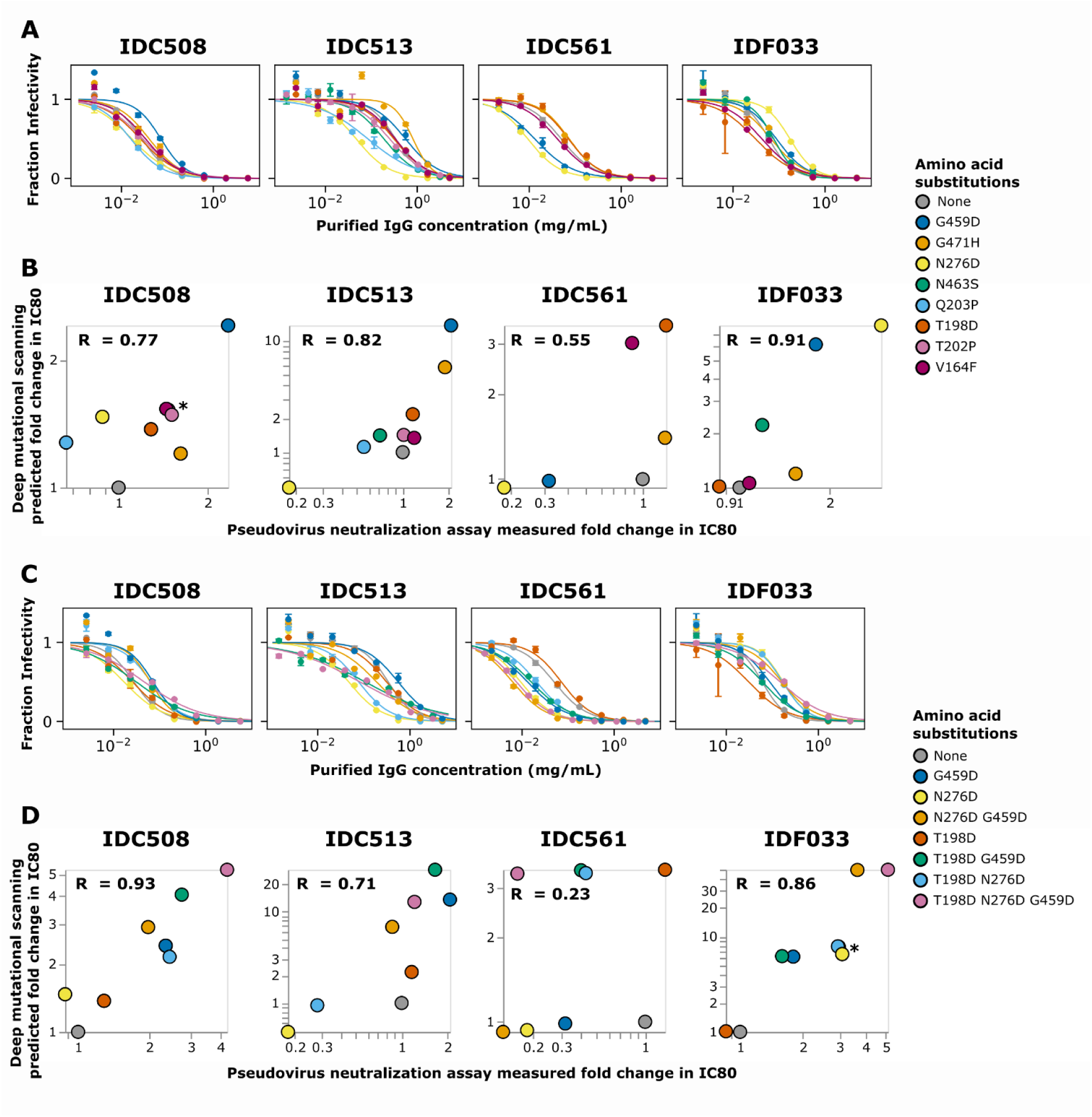
Pseudovirus neutralization assays to validate deep mutational scanning measurements. **(A)** Neutralization curves for unmutated BF520 Env and single mutants against purified IgGs from each sera. **(B)** Correlation of deep mutational scanning predicted fold change IC80s for single mutants versus fold change in IC80 measured in the neutralization assay. R indicates the Pearson correlation. (C) Neutralization curves for combinations of mutations (and the constituent individual mutations repeated from panel A). (D) Correlation of deep mutational scanning predicted fold change IC80s versus fold change IC80s measured in neutralization assays. Points in the scatter plots with asterisks overlap, so have been slightly jittered for clarity.

The correlation between the deep mutational scanning and neutralization assays was particularly good for strong escape mutations. For every serum, the tested mutations predicted to most strongly escape neutralization by the deep mutational scanning indeed increased the neutralization assays IC80 (Figure 7A,B). The correlation was less consistent for mutations that enhanced neutralization sensitivity rather than escape. For instance, N276D for serum IDC561 caused greater enhancement of neutralization sensitivity in the neutralization assays than predicted from the deep mutational scanning (Figure 7A,B). The reduced accuracy of the deep mutational scanning for identifying sensitizing mutations is probably because the mapping experiments were performed at relatively higher serum concentrations (typically exceeding the IC90 for unmutated BF520), making them better suited to identify escape rather than sensitizing mutations.

We also tested Env variants with combinations of mutations in neutralization assays, and again found a good correlation between the deep mutational scanning and neutralization assays (Figure 7C,D). For instance, the deep mutational scanning mapped IDC508 to have two epitopes, and pseudovirus neutralization assays using combinations of mutations supports this prediction (Figure 7C,D). Specifically, T198D is in one epitope of IDC508 while N276D and G459D are in the other epitope (Figure 6A)---and as predicted, N276D and G459D each cause more escape when combined with T198D (Figure 7C,D). Note, however, that the effects of these combinations of mutations is complex because N276D has some sensitizing effect on its own (Figure 7C,D). It could be that N276D simultaneously causes sensitization to some antibodies but escapes from others, which has been observed for other broad HIV neutralizing serum^7^. The greatest escape from IDC508 is caused by combining all three of T198D, N276D, and G459D, suggesting that this combination escapes a substantial fraction of the neutralizing antibodies in the serum (Figure 7C,D).

For sera IDC513 and IDC561, the deep mutational scanning predicted that combinations of mutations would not have substantially more escape than the single mutations with the highest effect, and this validated in neutralization assays (Figure 7C,D). Deep mutational scanning mapped IDC513 and IDC561 to each have one epitope (Figure 4A, Figure 5A). As expected for sera that target a single epitope, no combinations of mutations caused higher fold change IC80 in neutralization assays than the best escaping single mutations for IDC513 and IDC561 (Figure 7C,D). For IDC561, only one mutation tested in combinations (T198D) was measured in the deep mutational scanning to be an escape mutation, with the others being sensitizing mutations. Consistent with the deep mutational scanning, only the T198D single mutant caused escape in neutralization assays (Figure 7C,D), although similarly to the single mutants the sensitizing effects of N276D and G459D were underestimated by the deep mutational scanning. This poor estimation of the effects of sensitizing mutations leads to a worse overall correlation between the deep mutational scanning and validations for IDC561 (Figure 7C,D).

Despite being a sensitizing mutation for some sera, deep mutational scanning predicted mutations to site 276 to cause strong escape from serum IDF033 (Figure 6A), and as expected N276D caused a large increase in neutralization assay IC80 both alone (Figure 7A,B) and in combination with other mutations (Figure 7C,D). Consistent with the deep mutational scanning, combining N276D with another strong escape mutation, G459D, further increased the neutralization assay IC80 (Figure 7C,D).

## Discussion

We have described a lentiviral system for deep mutational scanning measurements of how mutations to Env affect antibody and serum neutralization. A major strength of this system is that it can measure the effects of combinations of mutations, enabling more effective mapping of escape from polyclonal serum that may target multiple epitopes. In addition, since the system is based on lentiviral vectors that can only undergo a single round of cellular infection, the experiments can be safely performed at biosafety level 2. Therefore, compared to prior Env deep mutational scanning that used fully replicative HIV^16, 18, 29, 35^, this system both enables more accurate measurements of how Env mutations affect neutralization and is safer and more convenient.

We have used the system to map the neutralization specificities to broadly neutralizing human anti-HIV sera. The mapping shows that although these sera all target Env’s CD4 binding site, they differ markedly in the actual epitopes that are the focus of the neutralizing response. Several of the sera have neutralization escape maps that strongly resemble those of characterized monoclonal antibodies. In one case, a serum resembles an antibody (1-18) actually isolated from the same individual. In another case, a serum resembles an antibody (3BNC117) from a different individual, suggesting convergent evolution of the same neutralizing antibody specificity across multiple individuals. However, one serum has an escape map best explained by neutralizing activity contributed by antibodies targeting two distinct epitopes near the CD4 binding site. Key predictions from all of these escape maps validated well in pseudovirus neutralization assays.

Neutralization by each of the sera we mapped is critically affected by the N276 glycan. However, the nature of this dependence is variable: for some sera neutralization is enhanced by disrupting the N276 glycan, where for others neutralization is escaped by disrupting this glycan. These findings are consistent with prior work. For instance, several described cases of broadly neutralizing antibody development have involved gain and loss of the N276 glycan during antibody-virus coevolution^7, 51^. Significant progress has been made in germline-targeting vaccinations that attempt to stimulate broadly neutralizing antibody precursors, but such strategies have not yet elicited responses in humans that neutralize heterologous viruses bearing the N276 glycan^52–54^. Here we observe strong effects of mutations at this glycan for several broad and potent sera targeting the CD4 binding site, suggesting the presentation of this glycan may be critical to any vaccination strategies that attempt to elicit such a response. In the future this method can be extended to identify mutations with sensitizing effects similar to N276 glycan knockout mutations, which could further aid sequential vaccine design efforts.

More broadly, the method described here can be used to map polyclonal neutralization escape in a variety of contexts. Combinations of broadly neutralizing antibodies targeting different regions of Env are being explored as potential therapies, and maps of escape from neutralization by single and combinations of antibodies could aid in antibody selection. Vaccine elicited sera can also be mapped to evaluate experimental vaccines and compare their neutralization activity to known broadly neutralizing antibodies or sera. Direct mapping of neutralization escape by polyclonal serum is therefore a useful tool for informing the design of both therapeutics and vaccines.

## Limitations of this study

Our experiments examined the effects of mutations in just one viral Env, BF520. However, mutations can have different effects across strains of HIV^29^, so these BF520 escape maps might not fully recapitulate escape mutations in other viral strains. For instance, BF520 is relatively resistant to neutralization by V3-specific antibodies^6^, which limits the ability to map escape from such antibodies using the current library. Future studies could extend the work we described here to Envs from more HIV strains to study these differences. Our study was also limited to a moderate number of sera capable of potently neutralizing BF520 Env. In addition, the method described here measures neutralization activity of sera and not other activities such as antibody binding, but it can be used in combination with other techniques like emPEM to study polyclonal serum antibody binding^13^.

## Acknowledgements

We thank Michael Emerman for scientific advice. We thank Dr. Michel Nussenzweig and Marina Caskey for a gift of 3BNC117 IgGs. This work was supported in part by the NIH/NIAID under grants R01AI141707, R01AI140891, and U01AI169385 to JDB. This work was supported in part by a Gates Foundation grant INV-004949 to JDB. CER was supported by the Viral Pathogenesis and Evolution training grant T32 AI083203. KHDC was supported by NIH/NIAID grant F30AI149928. BD was funded in part by EMBO under grant ALTF 81-2020. TCY was supported by the CMB training grant T32 GM007270 and the NSF graduate research fellowship DGE-2140004. JDB is an Investigator of the Howard Hughes Medical Institute. PS is supported by the Emmy Noether Program of the German Research Foundation (DFG, Project No: 495793173). This research was supported by the Genomics & Bioinformatics Shared Resource (RRID:SCR_022606), Flow Cytometry Shared Resource (RRID:SCR_022613), and Scientific Computing (NIH grants S10-OD-020069 and S10-OD-028685) of the Fred Hutch/University of Washington/Seattle Children’s Cancer Consortium (P30 CA015704).

## Competing Interests

JDB is on the scientific advisory boards of Apriori Bio, Invivyd, Aerium Therapeutics, and the Vaccine Company. JDB, KHDC, BD, ASD, and CER receive royalty payments as inventors on Fred Hutch licensed patents related to viral deep mutational scanning. ASD is currently an employee of Apriori Bio, though his contributions to this manuscript were performed when he was an employee of the Fred Hutch before he started work at Apriori Bio. A patent application encompassing aspects of this work has been filed by the University of Cologne and lists P.S. and F.K. as inventors. P.S., and F.K. received payments from the University of Cologne for licensed antibodies.

---

**Supplemental Figure 1:**
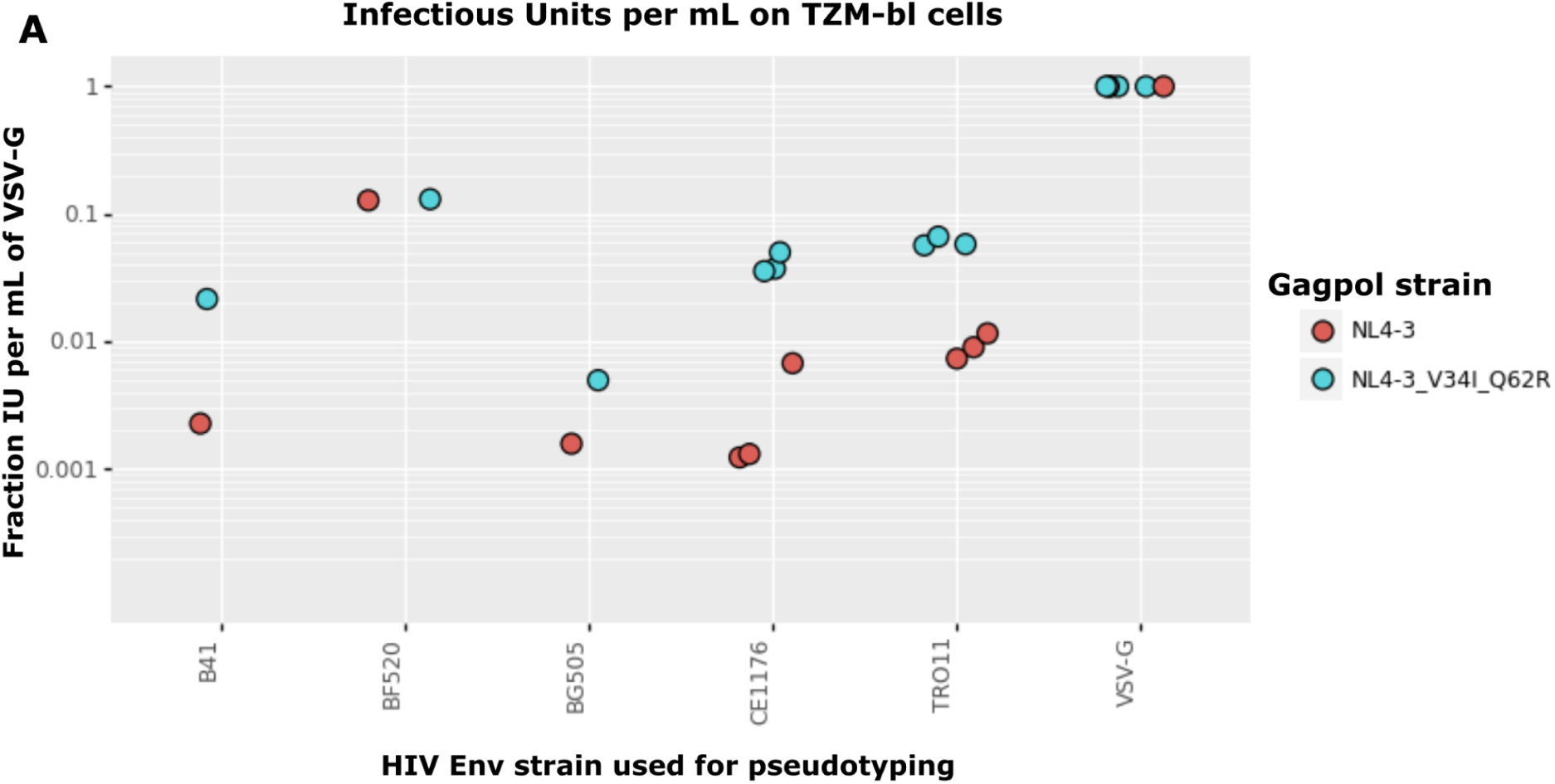
Titers of Env or VSV-G pseudotyped lentiviruses on TZM-bl cells. Lentiviruses were produced by transfecting 293T cells with the same ZsGreen reporter lentiviral backbone, Rev, and Tat expressing plasmids for each condition, along with an HIV Env from the indicated viral strain or a VSV-G expressing plasmid, and either a NL4-3 or NL4-3 V34I Q26R Gag-Pol expressing plasmid as indicated. Data are from different virus preparations and titering dates. The infectious units per mL on TZM-bl cells were normalized to VSV-G infectious units per mL by dividing each condition’s infectious units per mL by that of the VSV-G pseudotyped virus with the same Gag-Pol that was produced and titered on the same dates, performing this normalization to correct for batch effects. The titers for the VSV-G pseudotyped viruses ranged from ∼1.5-35 million infectious units per mL. Gag-Pol mutations V34I and Q62R were made based on previous studies that showed these mutations can rescue Env incorporation deficiencies^56, 57^.

**Supplemental Figure 2:**
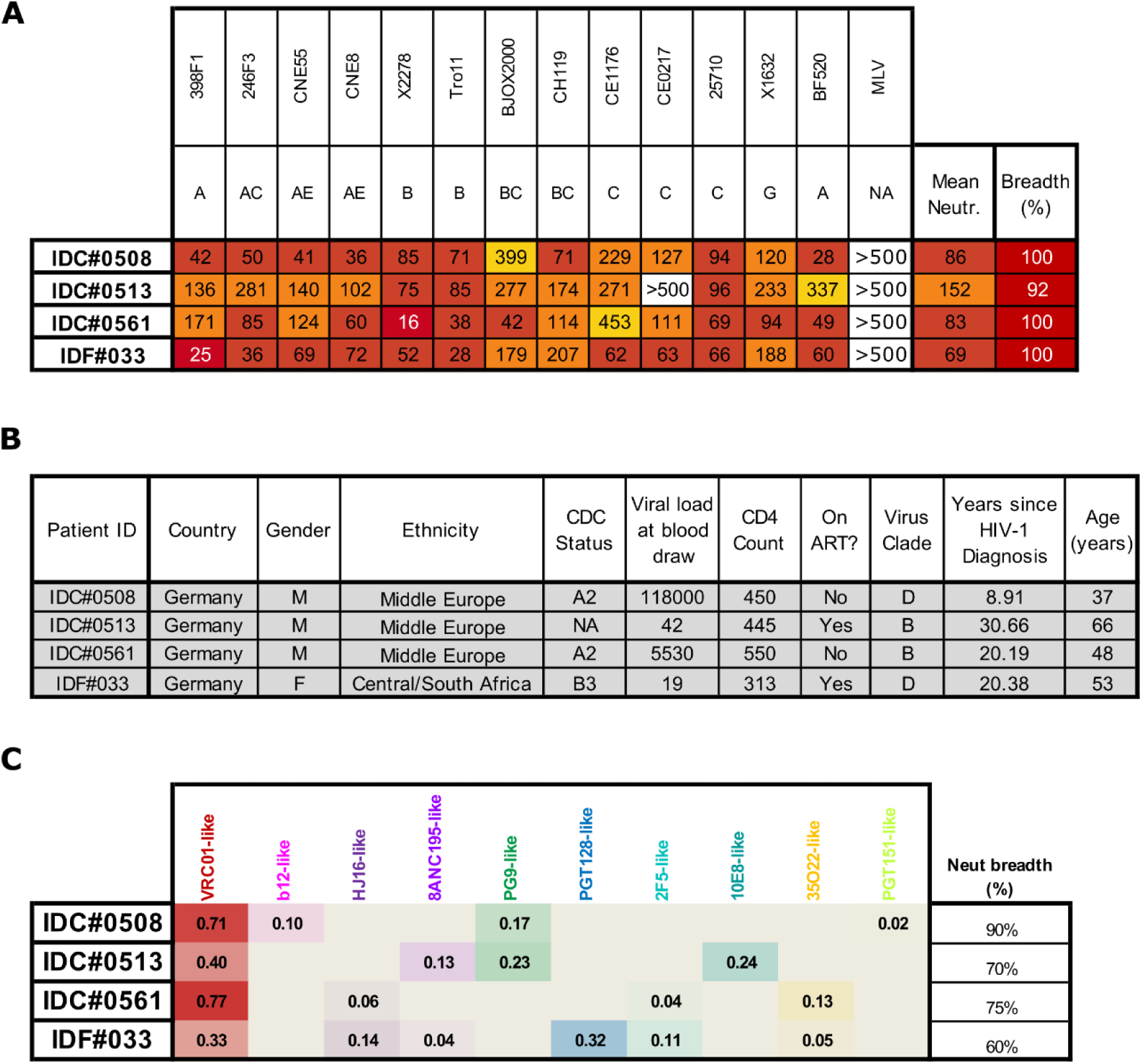
Broadly neutralizing human anti-HIV sera. (A) Neutralization of a global HIV panel by each serum^45^. Values reported are IC50 in ug/mL for purified IgGs. (B) Clinical data related to the individual from whom each serum was collected. (C) f61 neutralization fingerprinting results for each sera^14^.

**Supplemental Figure 3:**
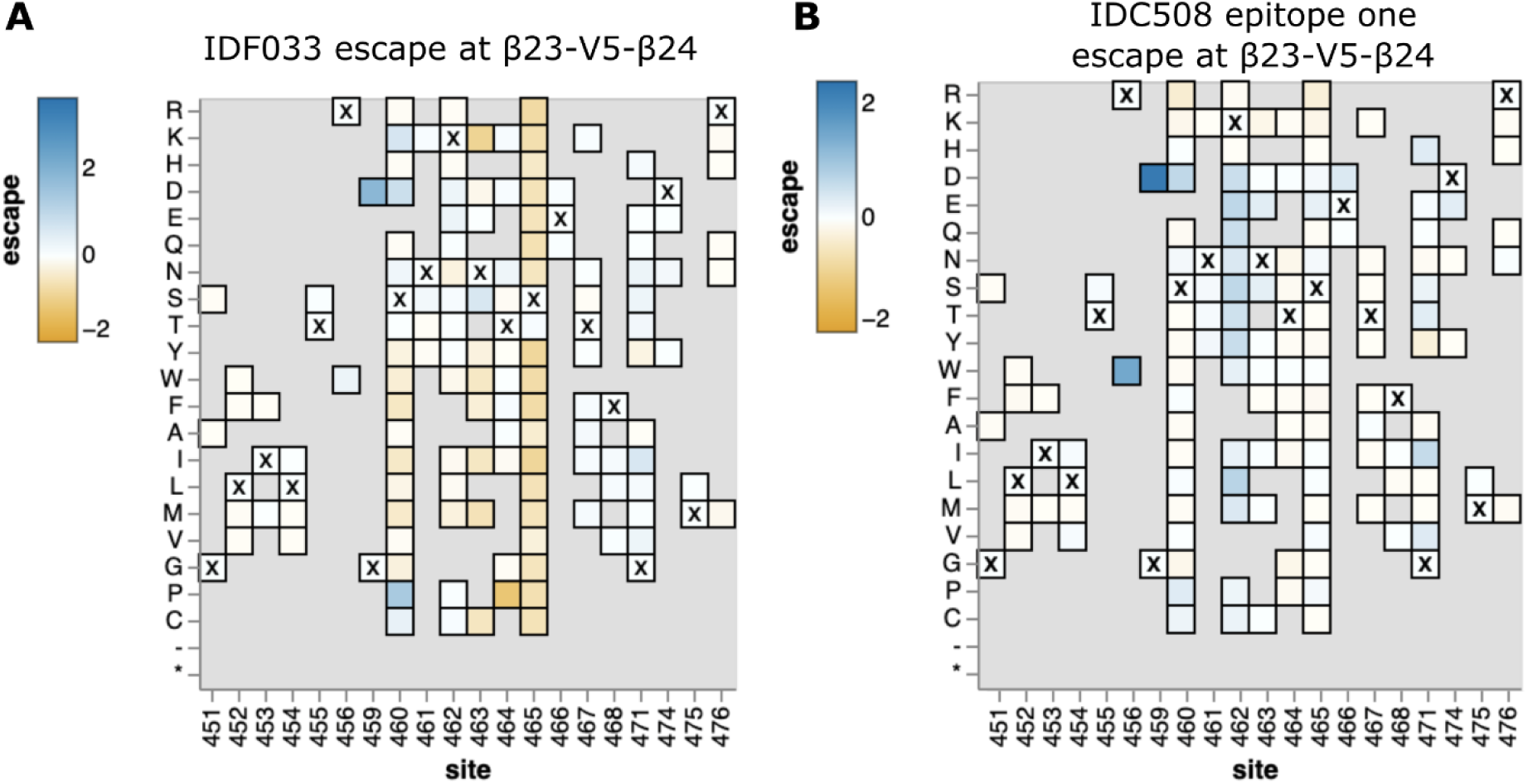
Zoomed in views of mutation-level escape at some key sites for IDF033 and IDC508. (A) Heatmap of escape of individual mutations in β23-V5-β24 for IDF033. Residues marked with X are wildtype residues in BF520. Residues grayed out are not present in the variant libraries. (B) Heatmap for IDC508 epitope 1. See https://dms-vep.github.io/HIV_Envelope_BF520_DMS_CD4bs_sera/IDF033_escape_plot.html and https://dms-vep.github.io/HIV_Envelope_BF520_DMS_CD4bs_sera/IDC508_escape_plot.html for interactive versions of the full Env escape maps for IDF033 and IDC508, respectively.

**Supplemental table 1:**
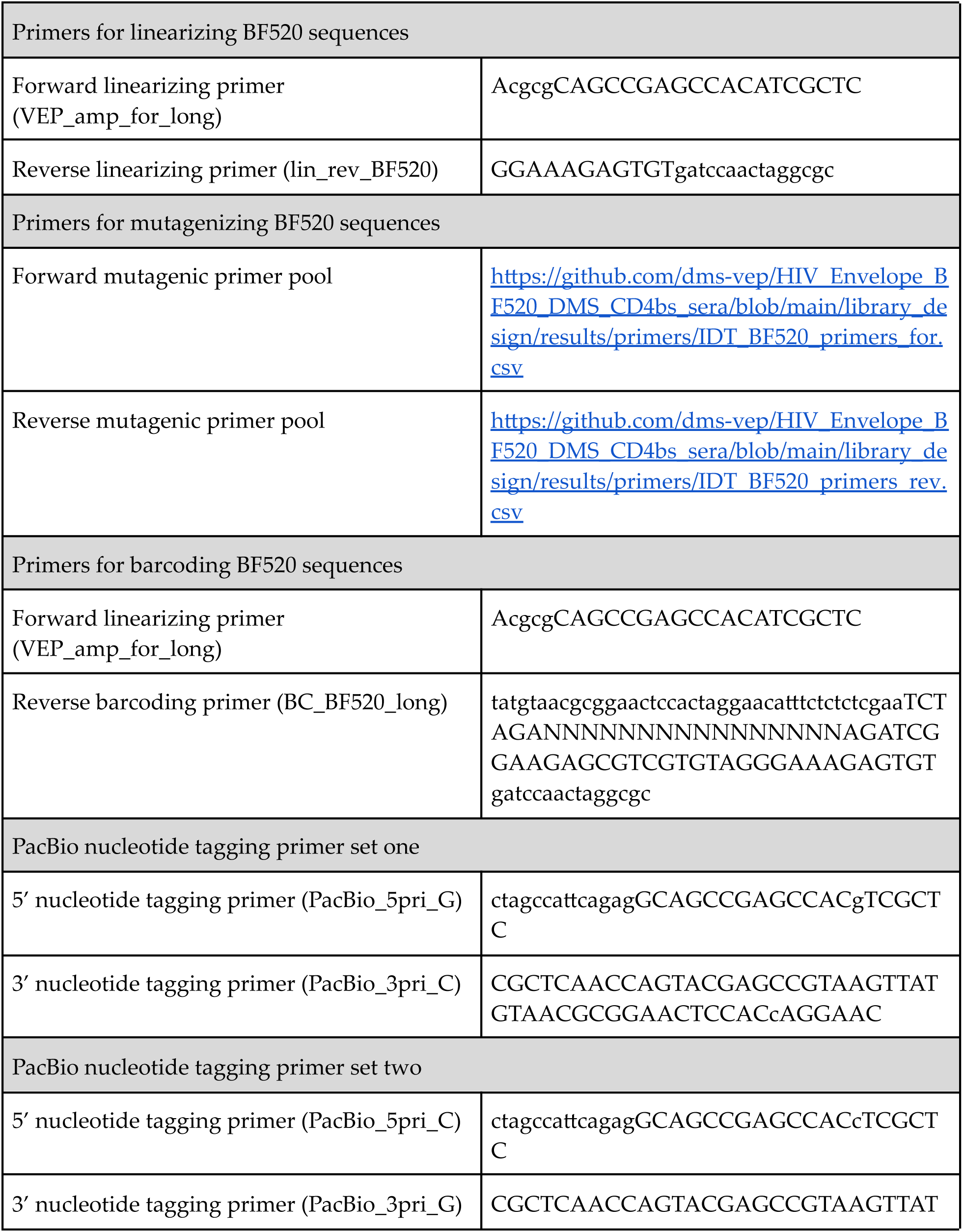
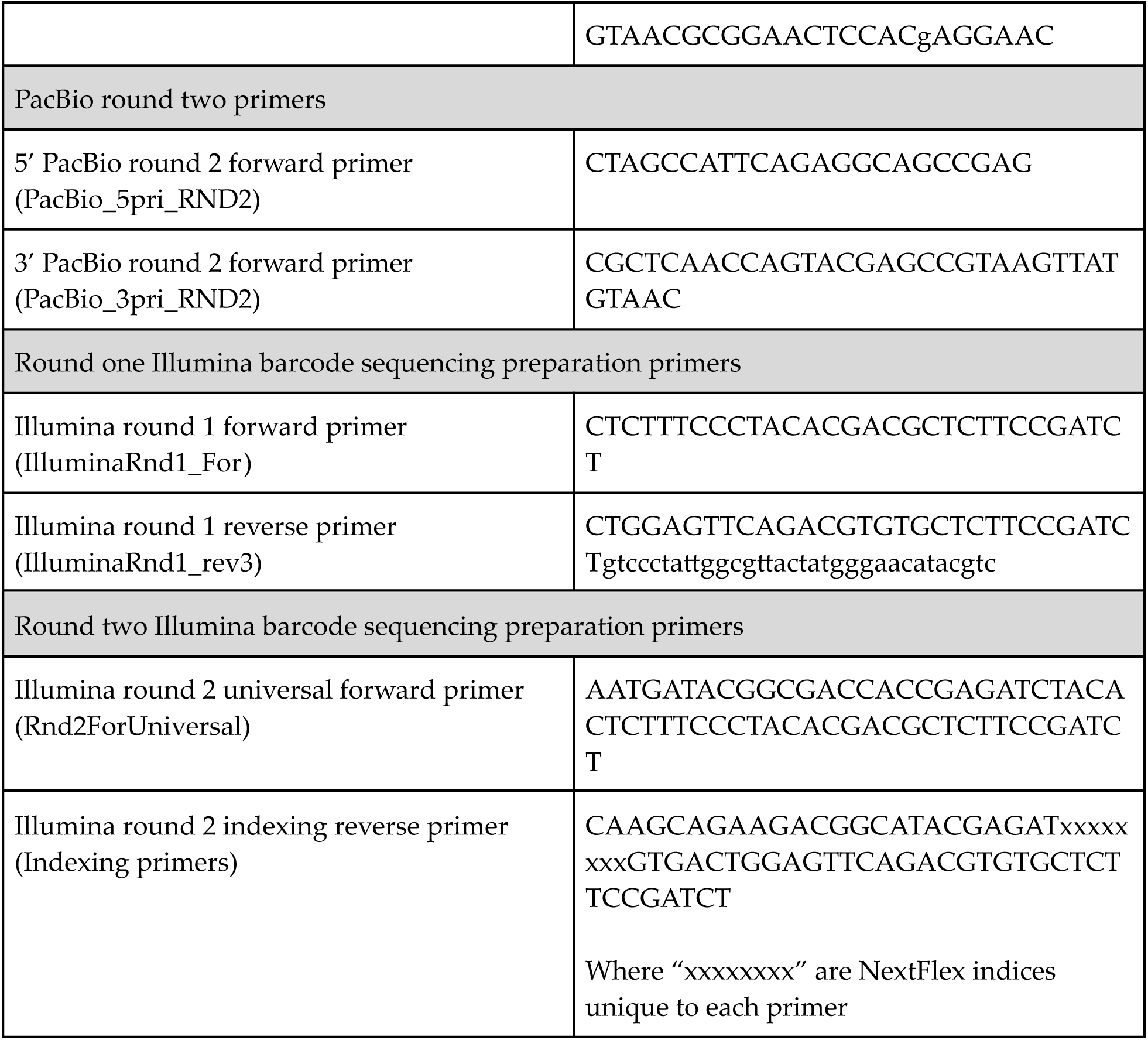
Primers and sequences

## Methods

### Design of lentivirus vector backbone

The lentivirus backbone we used is described in Dadonaite et al.^23^ See https://github.com/dms-vep/HIV_Envelope_BF520_DMS_CD4bs_sera/blob/main/plasmid_maps/lentivirus_backbone_plasmids/pH2rU3_ForInd_mCherry_CMV_ZsGT2APurR.gb for a map of the plasmid containing this backbone. Briefly, the backbone has a repaired 3’ LTR which allows it to be re-rescued after integrating into cells^23^, constitutive expression of ZsGreen and puromycin resistance as selectable markers for infection, and a TRE3G promoter that inducibly expresses HIV Env when the reverse tetracycline transactivator (rtTA) in the 293T-rtTA cells (described in Dadonaite et al.^23^ where they are referred to as HEK-293T-rtTAs) is induced by the presence of doxycycline. We used a codon optimized sequence of the HIV Env strain BF520.W14M.C2^26, 27^. See https://github.com/dms-vep/HIV_Envelope_BF520_DMS_CD4bs_sera/blob/main/plasmid_maps/lentivirus_backbone_plasmids/pH2rU3_ForInd_BF520.gb for a plasmid map containing the codon optimized BF520 sequence.

### Design of mutant libraries containing mostly functional mutants

To choose mutations to include in our mutant libraries based on prior BF520 deep mutational scanning^29^, we compared previously measured effects of all mutations vs the effects of stop codons. We retained mutations with an effect measured to be more positive than the 0.95 quantile of stop codon effect in the previous deep mutational scanning. We only retained three stop codons, at sites 100, 200, and 300, so we could use these as controls for selections.

We wanted to include mutations present in natural HIV sequences even if they had negative effects in previous deep mutational scanning, since these mutations are tolerable when combined with some other mutations and we wanted our neutralization selections to include most naturally occurring mutations. We downloaded the 2018 filtered web alignment of group M HIV-1 sequences without recombinants from the Los Alamos HIV sequence database^36^, and used it to identify any mutations relative to BF520 that were present more than once in the alignment, and retained these mutations for the mutant libraries in addition to those chosen above.

See https://github.com/dms-vep/HIV_Envelope_BF520_DMS_CD4bs_sera/tree/main/library_design for the analysis to choose these mutations. See https://github.com/dms-vep/HIV_Envelope_BF520_DMS_CD4bs_sera/blob/main/library_design/results/IDT_library_df.csv for the retained mutations.

### Design of primers for BF520 mutagenesis

See https://github.com/jbloomlab/TargetedTilingPrimers for the script used to generate primer sequences to make the chosen mutations. This script generates forward and reverse primers for each mutation which mutate that site to the most frequent codon of the desired mutant. Primer pools were ordered as oPools from Integrated DNA Technologies. See https://github.com/dms-vep/HIV_Envelope_BF520_DMS_CD4bs_sera/tree/main/library_design/results/primers for the primer sequences.

### Production of plasmids containing barcoded mutant BF520 sequences

See https://github.com/jbloomlab/CodonTilingPrimers for a general description of the PCR mutagenesis strategy we use here. The key difference is that we only ordered primers that introduced the targeted amino-acid mutations.

To mutagenize the BF520 sequences, a codon optimized BF520 sequence was first amplified from a plasmid containing the codon optimized BF520 sequence in a lentiviral backbone. See https://github.com/dms-vep/HIV_Envelope_BF520_DMS_CD4bs_sera/blob/main/plasmid_maps/lentivirus_backbone_plasmids/pH2rU3_ForInd_BF520.gb for the sequence of this plasmid. The PCR was performed with the following conditions: PCR mix: 18.5 µL H2O, 2.5 µL DMSO (to reduce off-target amplification), 1.5 µL 10µM forward linearizing primer (VEP_amp_for_long), 1.5 µL 10 µM reverse linearizing primer (lin_rev_BF520), 1 µL 10ng/µL BF520 template plasmid, and 25 µL 2x KOD Hot Start Master Mix (Sigma-Aldrich, Cat. No. 71842). Cycling conditions: (1) 95C/2min (2) 95C/20sec (3) 70C/1sec 54C/10sec, cooling at 0.5C/sec (5) 70C/40sec (6) Return to Step 2 x19.

The amplified, linearized BF520 sequence was gel purified using NucleoSpin Gel and PCR Clean-up kit (Takara, Cat. No. 740609.5) and then purified using Ampure XP beads (Beckman Coulter, Cat. No. A63881) at 1:1 sample to bead ratio.

The amplified BF520 sequence was then used in a modification of a previously described PCR mutagenesis technique^37^. Forward and reverse pools of codon tiling primers for generating specific mutations were generated using https://github.com/jbloomlab/TargetedTilingPrimers, as described above. In separate PCR reactions, the forward primer pool was used with the reverse linearizing primer and the reverse primer pool was used with the forward linearizing primer. The conditions for these PCR reactions were as follows: PCR mix: 7.7 µL H2O, 1.5 µL DMSO, 4 µL 3 ng/µL linearized BF520 template, 0.9 µL 10 µM forward or reverse primer pool, 0.9 µL reverse (lin_rev_BF520) or forward (VEP_amp_for_long) linearizing primer, and 15 µL 2x KOD Hot Start Master Mix. Cycling conditions: (1) 95C/2min (2) 95C/20sec (3) 70C/1sec (4) 50C/20sec, cooling at 0.5C/sec (50 70C/120sec (6) Return to Step 2 x9.

After the mutagenic PCRs, a joining PCR was performed using products from the forward and reverse primer pool mutagenic PCRs. The conditions for the joining PCRs were as follows: PCR mix: 4µL H2O, 4µL forward primer pool mutagenesis PCR product diluted 1:4 with H2O, 4µL reverse primer pool mutagenesis PCR product diluted 1:4 with H2O, 1.5µL 5µM forward linearizing primer (VEP_amp_for_long), 1.5µL 5µM reverse linearizing primer (lin_rev_BF520), and 15µL 2x KOD Hot Start Master Mix. Cycling conditions: (1) 95C/2min (2) 95C/20sec (3) 70C/1sec (4) 50C/20sec, cooling at 0.5C/sec (5) 70C/120sec (6) Return to Step 2 x19.

The resulting mutagenized BF520 sequences were gel purified and Ampure bead cleaned with a 1:1 product to beads ratio. These mutagenized sequences were then barcoded with random nucleotide barcodes using a PCR with the following conditions: PCR mix: 30 ng joining PCR product, 1.5µL 5µM forward linearizing primer (VEP_amp_for_long), 1.5µL 5µM reverse barcoding primer (BC_BF520_long), 15 µL 2x KOD Hot Start Master Mix, and fill to 30 µL with H2O. Cycling conditions: (1) 95C/2min (2) 95C/20sec (3) 70C/1sec (4) 50C/10sec, cooling at 0.5/sec (5) 70C/120sec (6) Return to Step 2 x9.

The barcoded mutagenized BF520 sequences were gel and Ampure bead purified, and then cloned into a lentiviral backbone containing plasmid as described in Dadonaite et al.^23^, with some modifications as follows. The barcoded mutagenized sequences were first cloned into an earlier version of the lentiviral backbone during system development. The map of the plasmid used can be found at https://github.com/dms-vep/HIV_Envelope_BF520_DMS_CD4bs_sera/blob/main/plasmid_maps/lentivirus_backbone_plasmids/pH2rU3_ForInd_mCherry_CMV_ZsG_NoBC_cloningvector.gb. The plasmid was digested with MluI and XbaI, and then gel and Ampure bead purified. The barcoded mutagenized BF520 sequences and the digested plasmid were eluted into H2O after Ampure bead purification, which we have observed results in higher Hifi assembly efficiency. We then used a 2:1 insert to vector ratio in a 1 hour Hifi assembly reaction using NEBuilder HiFi DNA Assembly kit (NEB, Cat. No. E2621). The Hifi assembly products were Ampure bead purified and eluted into 20 µL of H2O, which we have observed results in a higher electroporation efficiency. We used 2 µl of the purified HiFi product to transform 20 µl of 10-beta electrocompetent E. coli cells (NEB, C3020K). We performed 5 electroporation reactions for a final count of >5 million CFUs per library. We aimed for this high diversity of barcoded mutants in transformants to reduce the potential of barcode sharing in virus libraries, which we will describe in detail below. We plated the transformed cells on LB+ampicillin plates, incubated the plates overnight at 37 C, and scraped the plates the next day to collect the transformants. The OD600 of the collected bacteria were measured, and the bacteria were diluted to 15 OD600 and used in five separate 5 mL minipreps (QIAprep Spin Miniprep Kit, Cat. No. 27106X4) each, resulting in a total of ∼200 ug of plasmid being isolated for each replicate library. The rest of the bacteria were spun down in pellets and stored.

At a later stage of system development, we decided to move the barcoded mutagenized sequences into an improved version of the lentiviral backbone that uses puromycin selection rather than flow cytometry sorting to enrich infected cells when making the integrated mutant library cell lines. The map of this plasmid can be found at https://github.com/dms-vep/HIV_Envelope_BF520_DMS_CD4bs_sera/blob/main/plasmid_maps/lentivirus_backbone_plasmids/pH2rU3_ForInd_mCherry_CMV_ZsGT2APurR.gb. We decided to do this using restriction digest and ligation cloning of the library plasmids and the new lentiviral backbone plasmid. As an important note for future deep mutational scanning studies, this cloning strategy was not optimal. Since the barcoded mutagenized sequences were drawn from a plasmid pool with relatively limited diversity compared to mutagenic PCR products (a few million unique barcoded sequences vs >>billions of unique barcoded sequences), this cloning imposed an additional unintended bottleneck on the barcoded mutagenized sequence diversity. This meant that the final plasmid pools for each library had lower barcode diversity than intended, resulting in some degree of barcode sharing, described in a lower section. In the future, it is advised for similar deep mutational scanning strategies aiming for extremely high plasmid diversity to only clone from highly diverse mutagenic PCR products rather than any pre-existing mutant plasmid pool, which will always be limited in diversity by transformation efficiencies.

To move the barcoded mutagenized sequences into the improved lentiviral backbone, we digested each mutant plasmid pool and the new lentiviral backbone using MluI and XbaI. We gel extracted and Ampure bead cleaned the mutagenized barcoded inserts from the mutant plasmid pools and the cut lentiviral backbone vector, and eluted in Qiagen EB buffer (Cat. No. 19086). We then used T4 DNA ligase (New England BioLabs, Cat. No. M0202S) to ligate the inserts with the vector, using the following conditions: Reaction mix: 2 µL T4 DNA Ligase Buffer (10x), 50 ng Vector DNA, 45.35 ng insert DNA, 1 µL T4 DNA Ligase, and fill with H2O to 20 µL. We incubated the reaction at room temperature for 10 minutes, heat inactivated at 65C for 10 minutes, and then Ampure bead cleaned the product and eluted in 20 µL H2O. We then electroporated NEB 10beta cells (New England BioLabs, Cat. No. C3020K) following the protocol (http://www.neb.com/protocols/0001/01/01/electroporation-protocol-c3020) exactly. We performed five electroporations per library, for a total of ∼1 million CFUs per library. Again, as a note to future deep mutational scanning studies, the mutant plasmid pool restriction digest and ligation cloning strategy used here along with a transformation bottleneck <5 million CFUs is not recommended due to potential unintended bottlenecking of barcoded mutants.

### Production of cell lines storing BF520 mutant libraries

Production of cell line-stored BF520 mutant libraries was performed similarly to previously described in Dadonaite et al.^23^, with modifications (Figure 1B). This process involved the same steps of: 1) production of VSV-G pseudotyped lentiviruses carrying the barcoded mutant BF520 sequences, 2) infection of 293T-rtTA cells with the VSV-G pseudotyped viruses, and 3) selection for transduced cells using puromycin.

In order to not bottleneck the diversity of barcoded mutants at this step, we aimed to produce many more VSV-G pseudotyped viruses carrying the barcoded mutant BF520 sequences than the eventual desired library sizes of around 40,000 barcoded variants. We plated 500,000 293T cells per well in 6 well plates, and transfected 12 wells for each library. We used BioT (Bioland Scientific) for the transfections, and followed the manufacturer’s recommendations for the protocol and DNA / transfection reagent ratios. We transfected each well with 1 ug of lentiviral backbone plasmids carrying the barcoded mutagenized BF520 sequences, 250 ng of a HIV Tat expressing plasmid (HDM-tat1b), 250 ng of a HIV Rev expressing plasmid (pRC-CMV_Rev1b), 250 ng of a HIV Gag-Pol expressing plasmid (HDM-Hgpm2), and 250 ng of a VSV-G expressing plasmid (HDM_VSV_G). See https://github.com/dms-vep/HIV_Envelope_BF520_DMS_CD4bs_sera/tree/main/plasmid_maps for maps of these plasmids. We pooled the transfection supernatents for each library 48 hours post-transfection, filtered through a 0.45 µm SFCA syringe filter (Corning, Cat. No. 431220), and stored in 1 mL aliquots at −80C. We titrated these viruses based on the percent ZsGreen expression of cells infected with dilutions of virus as determined by flow cytometry, as described in Crawford et al.^58^ This yielded a total of >20 million viruses per library.

We used these VSV-G pseudotyped viruses to infect 293T-rtTA cells with approximately the same number of viruses as barcoded mutants that we desired in the final virus libraries. We aimed to avoid any bottlenecks in the barcoded mutant sequences before this step because recombination of pseudodiploid lentiviral genomes and mutations caused by lentiviral reverse transcription will alter barcode-mutant linkage during this step^23, 59–61^. We attempted to maintain high diversity in the barcoded sequences in prior steps to ensure each barcoded mutant-carrying lentiviral genome would have a unique barcode, so that barcodes would not be repeated in infected cells. After this step, each cell in the library storing cell lines will only have one integrated lentiviral genome with one barcoded mutant, so recombination in future steps is not an issue and mutations caused by reverse transcription in future steps will not alter mutant BF520 expression from these integrants and can be filtered in PacBio sequencing data, described below.

We aimed to infect the 293T-rtTA cells with between 30,000-40,000 variants per library. We first plated 500,000 293T-rtTA per well in ten six well plates. The next day, at the time of infection, we counted the cells per well in several wells. Based on the average count, we infected each well with the amount of infectious units required for a 0.005 multiplicity of infection, for five six well plates per library. Two days later, we determined the actual multiplicity of infection and infectious units per well for each library by determining the percent of infected cells by flow cytometry on ZsGreen expression and back-calculating the infectious units added per well based on that percentage and the average cell count per well at the time of infection. For each library we then pooled cells from the number of wells required for a total infectious units between 30,000-40,000. The pooled cells for each library were plated in a 10 cm plate.

Transduced cells were then selected for using puromycin selection, since infected cells expressed the puromycin resistance gene from the lentiviral genome while non-infected cells did not. Puromycin was added 24 hours after pooling at 0.75 ug/mL. 48 hours later, the cells were split into three 15 cm dishes per library with 0.75 ug/mL puromycin. 48 hours later, the media was replaced with fresh media plus 0.75 ug/mL puromycin. 48 hours later (a week after pooling), the cells for each library appeared all ZsGreen positive under a fluorescent microscope, and were expanded into one five layer flask (Falcon, Cat. No. 353144) per library. 24 hours later, half of the cells per library were frozen in 1 mL aliquots of 5 million cells in tetracycline-negative heat-inactivated fetal bovine serum (Gemini Bio, Cat. No. 100-800) with 10% DMSO, to be used in future virus library generation. The rest of the cells were used to generate mutant virus libraries as described below.

### Production of BF520 and VSV-G pseudotyped mutant virus libraries

Since each cell in the cell lines produced as described above contained one barcoded BF520 mutant, we were able to produce genotype-phenotype linked BF520 mutant virus libraries from them (Figure 1B). We did this by plating 100 million cells per flask in two five layer flasks per library in 150 mL of tetracycline free D10. 24 hours later, we transfected each flask using BioT by using 225 µL of BioT mixed with 7.5 mL of DMEM and a DNA mix containing 50 ug of each lentivirus helper plasmid (Tat, Rev, and Gag-Pol). We also induced Env expression at the time of transfection by adding doxycycline to a final concentration of 100 ng/mL. 48 hours later, the supernatant for each library was filtered through a 0.45 µM SFCA filter (Nalgene, Cat. No. 09-740-44B). The filtered virus was then concentrated using ultracentrifugation with a 20% sucrose cushion at 100,000 g for one hour. The viruses were resuspended in 500 µL of DMEM, and were typically around ten million infectious units per mL. We then stored these viruses at −80C.

We also generated VSV-G pseudotyped viruses from the library cell lines to use for PacBio sequencing and as controls for selections on the effects of mutations on BF520 function, described below. We did this by plating four million cells per plate in three 10 cm dishes for each library, and transfecting each plate 24 hours later using BioT according to the manufacturer’s recommendations. For the DNA mix, we used 2.5 ug of each lentivirus helper plasmid (Tat, Rev, and Gag-Pol) and a VSV-G expressing plasmid (four plasmids, 10 ug total DNA) per plate. 48 hours later we pooled the supernatants for each library and filtered them through a 0.45 µM SFCA filter. We then stored these viruses at −80C.

### PacBio sequencing of mutants present in mutant libraries

We used long-read sequencing PacBio sequencing to simultaneously determine the composition of the mutant libraries contained in the library cell lines and link mutants with their random nucleotide barcodes. First, we plated 1 million 293Ts per well in poly-L-lysine coated six well plates (Corning, Cat. No. 356515). 24 hours later, we infected two wells of cells with 1 million infectious units of +VSV-G library virus per well, for each library. Six hours later, we removed the media, washed the cells with PBS, and miniprepped each well, which isolates unintegrated lentivirus genomes as described previously.^23, 35^ Each well was miniprepped independently and eluted using 50 µL of EB.

A two-step PCR strategy was then used to amplify the barcoded mutant BF520 sequences for PacBio sequencing, as described previously.^23^ Briefly, the miniprepped products for each library were split into two short-cycle initial PCRs that attached single nucleotide tags to each end of the amplicon that were unique for each PCR. The products of these initial PCRs were then pooled for each library for longer cycle PCRs to amplify enough DNA for PacBio sequencing. The single nucleotide tags from the initial PCRs then allowed us to later estimate the amount of strand exchange that occurred in the longer cycle PCRs based on the frequency of tags found together in PacBio sequences that were from different first round PCRs. The first round PCR is a low cycle number to minimize the probability of strand exchange during it, and the number of cycles in the second PCR was lowered as much as possible to minimize strand exchange while still generating enough DNA for PacBio sequencing. Here are the conditions used for the first round of PCRs: PCR mix: 10 µL of miniprep product, 1 µL of 10 µM 5’ nucleotide tagging primer (PacBio_5pri_G or PacBio_5pri_C), 1 µL of 10 µM 3’ nucleotide tagging primer (PacBio_3pri_C or PacBio_3pri_G), 20 µL KOD Hot Start Master Mix, and 8 µL H2O. Cycling conditions: (1) 95C/2min (2) 95C/20sec (3) 70C/1sec (4) 60C/10sec, cooling at 0.5/sec (5) 70C/60sec (6) Return to Step 2 x7 (7) 70C/60sec.

The PCR products were cleaned with Ampure beads with a 1:1 product to beads ratio and eluted into 35 µL of EB. We then used the following conditions for the second round of PCRs: PCR mix: 10.5 µL of first variant tag set round 1 PCR product, 10.5 µL of second variant tag set round 1 PCR product, 1 µL of 10 µM 5’ PacBio round 2 forward primer (PacBio_5pri_RND2), 1 µL of 10 µM 3’ PacBio round 2 reverse primer (PacBio_3pri_RND2), and 25 µL KOD Hot Start Master Mix. Cycling conditions: (1) 95C/2min (2) 95C/20sec (3) 70C/1sec (4) 60C/10sec, cooling at 0.5/sec (5) 70C/60sec (6) Return to Step 2 x10 (7) 70C/60sec. The PCR products were Ampure bead cleaned, and each eluted into 40 µL of EB. The cleaned products for each library were pooled. Each library pool was then barcoded for PacBio sequencing using SMRTbell prep kit 3.0, bound to polymerase using Sequel II Binding Kit 3.2, and then sequenced using a PacBio Sequel IIe sequencer with a 20-hour movie collection time. The data were analyzed as described below (section “PacBio sequencing data analysis”).

### Barcode amplification for Illumina sequencing of mutants after selections

After the above step using PacBio sequencing to link each mutant and barcode, future experimental steps only require short read sequencing of barcodes to determine changes in variant frequencies across conditions. We amplified barcodes for sequencing as previously described in Dadonaite et al.^23^ with slight modifications, repeated here. A first round of PCR was used to amplify the barcodes using a forward primer that aligns to the Illumina Truseq Read 1 sequence upstream of the barcode in our lentiviral backbone and a reverse primer that annealed downstream of the barcode and overlapped with the Illumina Truseq Read 2 sequence. This PCR used the following conditions: PCR mix: 22 µL of miniprepped selection sample, 1.5 µL of 10 µM 5’ Illumina round 1 forward primer (IlluminaRnd1_For), 1.5 µL of 10 µM 3’ Illumina round 1 reverse primer (IlluminaRnd1_rev3), and 25 µL KOD Hot Start Master Mix. Cycling conditions: (1) 95C/2min (2) 95C/20sec (3) 70C/1sec (4) 58C/10sec, cooling at 0.5/sec 70C/20sec (6) Return to Step 2 x27.

The PCR products were Ampure bead cleaned with a 1:3 product to beads ratio, and then DNA concentration was quantified using a Qubit Fluorometer (ThermoFisher). A second round of PCR was then performed using a forward primer that annealed to the Illumina Truseq Read 1 sequence and had a P5 Illumina adapter overhang, and reverse primers from the PerkinElmer NextFlex DNA Barcode adaptor set that annealed to the Truseq Read 2 site and had the P7 Illumina adapter and i7 sample index. This PCR used the following conditions: PCR mix: 20 ng of round 1 product as determined by Qubit, 2 µL of 10 µM 5’ Illumina round 2 universal forward primer (Rnd2ForUniversal), 2 µL of 10 µM 3’ Illumina round 2 indexing reverse primer (Indexing primers), 25 µL KOD Hot Start Master Mix, and fill to 50 µL total using H2O. Cycling conditions: (1) 95C/2min (2) 95C/20sec (3) 70C/1sec (4) 58C/10sec, cooling at 0.5/sec (5) 70C/20sec (6) Return to Step 2 x19.

The DNA concentration of each round 2 PCR product was quantified using Qubit. The samples were pooled at an even ratio, gel purified and Ampure bead cleaned at a 1:3 sample to beads ratio, and then sequenced using either P2 or P3 reagent kits on a NextSeq 2000. The data were analyzed as described below (section “Illumina barcode sequencing data analysis”).

### Selections on effects of mutations on the function of BF520

To measure the effects of mutation on BF520 mediated entry into cells, we infected cells with VSV-G and non-VSV-G pseudotyped mutant virus libraries separately. To do this, we plated 1 million TZM-bl cells in each well of six well plates. 24 hours later, we infected each well with ∼1 million infectious units of VSV-G or non-VSV-G pseudotyped mutant virus depending on the condition. We used this amount of virus because we aimed to use >20x the size of each mutant library during infections, so that each barcoded mutant would be present more than once and less likely to be randomly bottlenecked during the selections. During infections, we added 100 ug/mL DEAE dextran, which improves the infectivity of Env pseudotyped viruses and results in less random bottlenecking of mutants during infections.^16, 35^ 12 hours after infection, the cells were washed with PBS, miniprepped using a QIAprep Spin Miniprep Kit to isolate unintegrated lentivirus genomes as described previously^23, 35^, and eluted into 30 µL of EB. To improve the DNA recovery, the EB was run through the column twice, incubating at 55C for five minutes before spinning each time. The eluent was then used in the barcode sequencing prep described above.

### Production of VSV-G pseudotyped standard viruses for neutralization selections

For each selection using antibodies or sera, we spiked in a small amount of a separately produced only-VSV-G pseudotyped virus pool carrying known barcodes to act as neutralization standards by enabling conversion of barcode counts to absolute neutralization values (See Figure 3 of Dadonaite et al.^23^). We produced these viruses exactly as described in Dadonaite et al.^23^ Briefly, 293T-rtTA cells were transduced at a low multiplicity of infection with a pool of lentiviruses carrying a small set of known barcodes but no viral entry protein in their genomes. Transduced cells were selected for using flow cytometry cell sorting on ZsGreen expression, and then standard viruses were produced by transfecting the cells with the lentiviral helper plasmids and a plasmid expressing VSV-G. The result of this process was a standard virus pool with known barcodes that was produced in the same manner as mutant libraries but did not contain any viral entry protein mutants.

### Selections on effects of mutations on neutralization escape

We aimed to perform antibody and serum selections at concentrations between the IC90-IC99.9 for each antibody and serum. We used a spread of concentrations in this range because it is difficult to estimate IC9X concentrations and we wanted to use a spread of high neutralization levels to fit our biophysical escape models.^22^ When performing selections using antibodies or serum with the mutant virus libraries, we spiked-in the VSV-G pseudotyped neutralization standard viruses to be 0.5-1% of the total infectious units in the virus pool. From this combined virus pool, 1 million infectious units per selection were incubated with antibody or serum at the desired concentration for one hour. After the incubation, the volume of each condition was raised to 2 mL with 100 ug/mL DEAE dextran using D10 with the appropriate amount of DEAE dextran. Each condition was used to infect one well of TZM-bl cells in a six well dish plated at 1 million cells per well 24 hours prior. 12 hours after infection, the cells were washed with PBS, miniprepped, and eluted into 30 µL of EB. To improve the DNA recovery, the EB was run through the column twice, incubating at 55C for five minutes before spinning each time. The eluent was then used in the barcode sequencing prep described above.

### Validation pseudovirus neutralization assays

Plasmids containing BF520 with mutations used in pseudovirus neutralization assays were ordered from Twist in the HDM plasmid (https://github.com/dms-vep/HIV_Envelope_BF520_DMS_CD4bs_sera/blob/main/plasmid_maps/viral_entry_protein_expression_plasmids/HDM_BF520.gb). To produce viruses pseudotyped with each BF520 mutant, we first plated 500,000 293T cells per well in six well plates. 24 hours later we transfected 1 ug of a ZsGreen and Luciferase expressing lentivirus backbone plasmid (https://github.com/dms-vep/HIV_Envelope_BF520_DMS_CD4bs_sera/blob/main/plasmid_maps/lentivirus_backbone_plasmids/pHAGE6-wtCMV-Luc2-BrCr1-ZsGreen-W-1247.gb), 250 ng of each lentiviral helper plasmid (Tat, Rev, and Gagpol), and 250 ng of the HDM plasmid expressing the desired BF520 mutant into each well. We collected the viruses 48 hours later by filtering the supernatant through a 0.45 µm SFCA syringe filter and storing the virus at −80C.

To titrate these viruses for use in neutralization assays, we first plated 25,000 TZM-bl cells per well in clear bottom, poly-L-lysine coated, black walled 96 well plates (Greiner, Cat. No. 655930). 24 hours later, we serially diluted each mutant BF520 pseudotyped virus and infected the cells. 48 hours after infection, we used the Bright-Glo Luciferase Assay System (Promega, E2610) to measure relative light units (RLUs) for each dilution. We estimated the average RLU/µL for each BF520 mutant within a linear range based on its dilution curve. Note, this method and the following described neutralization assay are not the same as a typical TZM-bl neutralization assay, since Luciferase expression will be driven from the lentiviral genome of the infecting virus rather than the pre-integrated Tat-driven Luciferase in the TZM-bl cells, as there is will be no Tat expressed from these lentiviruses. We chose to do this rather than using ΔEnv HIV pseudoviruses in typical TZM-bl neutralization assays so that there was no chance of the BF520 Env mutants with combinations of escape mutations to CD4 binding site antibodies or sera recombining into full-length replicative HIV.

For neutralization assays, we plated 25,000 TZM-bl cells per well in clear bottom, poly-L-lysine coated, black walled 96 well plates. 24 hours later, we serially diluted each antibody or sera, and then incubated each dilution with each mutant BF520 pseudotyped virus for one hour. We then added an equal volume of D10 with DEAE dextran to a final DEAE dextran concentration of 100ug/mL, and infected the TZM-bls. 48 hours later, we used the Bright-Glo Luciferase Assay System to measure RLUs for each dilution.

To calculate fraction infectivity, we subtracted the average background reading of RLUs from uninfected cells from each condition, and then divided the RLU of each antibody or serum dilution by the average RLUs from cells infected by virus that was incubated with media rather than antibodies or sera. The fraction infectivities were used to fit neutralization curves using *neutcurve* (https://jbloomlab.github.io/neutcurve/). We compared fold change IC80 rather than IC50 for our interpretation of the neutralization assays because our deep mutational scanning selections were performed at high levels of neutralization (>IC90 for wildtype BF520).

### Experimental replicates

We performed one or two replicates of each selection with two independent mutant libraries for each experiment. See https://dms-vep.github.io/HIV_Envelope_BF520_DMS_CD4bs_sera/avg_muteffects.html for correlation plots of functional effects of mutations across replicates and see https://dms-vep.github.io/HIV_Envelope_BF520_DMS_CD4bs_sera/avg_antibody_escape.html for correlation plots of escape effects of mutations across replicates for each antibody and serum. Throughout the paper we report the median across these replicates.

### Cell lines

HEK-293T cells were from ATCC (CRL3216), TZM-bl cells were from the NIH AIDS Reagent Program (ARP-8129, contributed by Dr. John C. Kappes, Dr. Xiaoyun Wu and Tranzyme Inc.), and 293T-rtTA expressing cells were produced as previously described in Dadonaite et al.^23^ where they are referred to as HEK-293T-rtTAs. All cell lines were grown in D10 media (Dulbecco’s Modified Eagle Medium with 10% heat-inactivated fetal bovine serum, 2 mM l-glutamine, 100 U/mL penicillin, and 100 µg/mL streptomycin). To avoid rtTA activation and mutant BF520 expression earlier than intended, 293T-rtTA cells were grown in D10 made with tetracycline-free fetal bovine serum (Gemini Bio, Cat. No. 100-800).

### Antibodies

3BNC117 IgGs were a gift from Dr. Michel Nussenzweig and Dr. Marina Caskey, and PGT151 and 1-18 IgGs were produced by Genscript based on publicly available sequences.

### Patient plasma samples and IgG isolation

Blood samples were obtained under protocols approved by the Institutional Review Board (IRB) of the University of Cologne (protocols 13-364 and 16-054) and the local IRBs and all participants provided written informed consent. HIV-1 infected patients were recruited at private practices and/or hospitals in Germany. Plasma samples were obtained and stored at −80°C until further use. Prior to IgG isolation, plasma samples were heat-inactivated at 56°C for 40 minutes. IgGs were isolated through an overnight incubation with Protein G Sepharose (GE Life Sciences) at 4°C, followed by elution with 0.1 M glycine (pH=3.0) using chromatography columns. The eluted IgGs were buffered in 1 M Tris (pH=8.0) and then underwent buffer exchange to phosphate-buffered saline (PBS) and concentration using Amicon 30 kDa spin membranes (Millipore). The purified IgGs were stored at 4°C until further use.

## Computational Methods

### Computational pipeline overview

For analyzing deep mutational scanning of viral entry protein, we use a common, modular pipeline. See https://github.com/dms-vep/dms-vep-pipeline for this pipeline. For this paper, we used version 2.0.1 of *dms-vep*-*pipeline*. We created a repository for the analyses performed in this paper. See https://github.com/dms-vep/HIV_Envelope_BF520_DMS_CD4bs_sera for this repository. This repository includes the main *dms-vep*-*pipeline* as well as all of the scripts, notebooks, and settings necessary to recreate the analysis. Some key results files can be found in this repository, but some results files that are too large are not tracked in the online repository. The pipeline also produces HTML rendering of the key analyses and interactive plots. See https://dms-vep.github.io/HIV_Envelope_BF520_DMS_CD4bs_sera/ for these pages. These pages are the best way to explore the analyses and interactive plots of the results.

### PacBio sequencing data analysis

We used *alignparse* (see https://jbloomlab.github.io/alignparse/ for documentation) to analyze the PacBio sequencing data^62^. The PacBio CCSs went through several filtering steps before we determined which BF520 mutants were linked to which barcodes. First, we looked for evidence of strand exchange during the PacBio sequencing prep PCRs by computing the fraction of CCSs that contained unexpected pairs of single nucleotide tags, such as pairs of nucleotide tags from different round one PCRs or any wildtype nucleotides. These sequences represented just <1% of the CCSs, and were filtered out.

Next, we computed empirical accuracies for each CCS, which represent the fraction of CCSs with the same barcode that report the same BF520 sequence. The empirical accuracies were around 0.60, slightly less than our previously described SARS-2 spike libraries^23^. Inaccuracies in the BF520 sequences are due to a combination of factors including reverse transcription errors, sequencing errors, strand exchange during PCR, and, importantly, actual linkage of the same barcode sequence with two or more BF520 mutants due to having the same barcode with different BF520 mutants integrated into different cells. As noted in above sections, we know that we unintentionally bottlenecked the barcoded BF520 mutants to a lower level than desired, which likely resulted in some barcode sharing between variants and contributed to the reduction in empirical accuracies.

We filtered out any barcodes that had less than three CCSs or had minor fractions of substitutions or indels above 0.4. This removes consensus sequences that we are not confident in due to not having enough CCSs for that barcode or evidence of multiple BF520 mutants sharing that barcode. The remaining consensus sequences of barcoded mutant BF520 sequences were then saved in barcode / variant lookup table files. See https://dms-vep.github.io/HIV_Envelope_BF520_DMS_CD4bs_sera/build_pacbio_consensus.html for the analysis described in this section and https://github.com/dms-vep/HIV_Envelope_BF520_DMS_CD4bs_sera/blob/main/results/variants/codon_variants.csv for the barcode / variant lookup tables.

### Illumina barcode sequencing data analysis

We used the parser found at https://jbloomlab.github.io/dms_variants/dms_variants.illuminabarcodeparser.html/ to determine the counts of each variant in each selection condition. For each experiment, barcoded mutants were only retained if their “pre-selection” counts, such as counts in no-antibody conditions for antibody selections or VSV-G condition counts for functional selections, were above thresholds specified in the configuration file for the analysis. See https://github.com/dms-vep/HIV_Envelope_BF520_DMS_CD4bs_sera/blob/main/config.yaml for the configuration file. This was done because these barcoded mutants are more likely to be randomly bottlenecked during the infection step of selections and to have highly noisy scores due to the computation of functional and antibody escape scores (described below).

### Modeling the effects of mutations on Env function

We modeled the functional effects of mutations on BF520 as described previously in Dadonaite et al.^23^ Briefly, we computed functional scores for each barcoded mutant *m* defined as 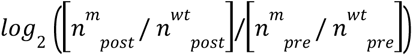 where where *n^m^* _*post*_ is the count of mutant *m* in the post-selection (BF520-pseudotyped) infection, *n^m^ _pre_* is the count of mutant *m* in the pre-selection (VSV-G-pseudotyped) infection, and *n^wt^* _*post*_ and *n^wt^* _*post*_ are the counts of all unmutated (wildtype) barcoded variants in each condition. We then used global epistasis models^41, 42^ to deconvolve the functional scores of these thousands of mutants with combinations of mutations into estimates of the effects of each individual mutation. Under these models, each mutation has a “latent effect” on a “latent phenotype” defined as an unmeasured phenotype where mutations interact additively. These latent effects and the latent phenotype score for a mutant are then transformed to the actually measured “observed phenotype” through a nonlinear function. This process results in more accurate estimates for the effects of mutations by modeling some of the epistasis between mutations through the nonlinear relationship between the latent and observed phenotypes. See https://dms-vep.github.io/HIV_Envelope_BF520_DMS_CD4bs_sera/fit_globalepistasis.html for the models, see https://dms-vep.github.io/HIV_Envelope_BF520_DMS_CD4bs_sera/muteffects_latent_heatmap.html for the latent effects of mutations averaged across selections, and see https://dms-vep.github.io/HIV_Envelope_BF520_DMS_CD4bs_sera/muteffects_observed_heatmap.html for the observed effects of mutations averaged across selections. For Figure 2C, we used the observed effects of mutations, since these are the most accurate representation of the effects of mutations on BF520.

### Modeling the effects of mutations on antibody and serum escape

We modeled the effects of mutations on antibody and serum escape as described previously in Dadonaite et al.^23^ Briefly, we calculated the non-neutralized fraction of each variant at each antibody calculation to get the probability of escape of each variant at each concentration. We then used the software package *polyclonal*^22^ version 4.1 (see https://jbloomlab.github.io/polyclonal/ for documentation) to estimate the effects of each individual mutation on escape using a biophysical model. Under this model, antibodies, mixtures of antibodies, or polyclonal serum can target Env at one or more epitopes. Measured probability of escape of each variant is modeled as a result of how the mutations it has escape the neutralization activity of each epitope. The measured probability of escape of combinations of mutations across mutants is used to optimize the number and sites of antibody epitopes and give each mutation escape scores corresponding to its contributions to escape for each epitope when present. Mutations that have no effect on escape will have scores of zero, while mutations that cause escape will have scores >0. The summed escape scores for each site are the y-axis values displayed in the line plots in each figure and used to color the PDB structures seen in each figure. The individual escape scores for each mutation can be seen in the heatmaps of the linked interactive plots, like the ones seen in Figure 3B and Supplemental Figure 3.

The models are also able to predict arbitrary inhibitory concentrations for Env mutants, such as an IC50 or IC80 for serum IDC508 against BF520 with mutations T198D and N276D. This is done by determining the effect of each mutation on escape from each epitope in the neutralizing activity of the serum, and then predicting the non-neutralized fraction of virus depending on the degree each epitope’s activity is escaped and the contribution of each epitope to the total neutralizing activity^22^. These predictions were generated for the BF520 mutants used in the neutralization assays depicted in Figure 3D and Figure 7. We chose to compare the fold change in IC80s because these values are similar to the level of neutralization we used in our deep mutational scanning selections.

We constrained the model for each antibody to have one epitope, while sera could have up to two epitopes. For figures in this manuscript and the interactive figures, we filter the mutations in the default view by requiring mutations to be present in at least three unique variants and to have a functional effect above −1.5. We filter mutations with low functional scores because variants with these mutations typically have low counts in no-antibody selections, which can cause high amounts of noise in their probability of escape scores. See https://dms-vep.github.io/HIV_Envelope_BF520_DMS_CD4bs_sera/ for interactive plots, notebooks detailing the fitting of these models, and PDBs with b-factors containing the escape values for each model.

## Resource Availability

### Lead Contact

Further information and requests for reagents and resources should be directed to and will be fulfilled by the Lead Contact, Jesse Bloom.

### Materials Availability

BF520 Env mutant libraries generated in this study will be made available on request by the Lead Contact with a completed Materials Transfer Agreement.

### Data and code availability

All data is available via links provided in the Methods section and Key Resources table. See https://github.com/dms-vep/HIV_Envelope_BF520_DMS_CD4bs_sera for the full analysis pipeline and key results. The raw sequencing data for this study can be found in the NCBI Sequence Read Archive under BioProject number PRJNA947170.

## Notes

https://dms-vep.github.io/HIV_Envelope_BF520_DMS_CD4bs_sera/

